# A tale of two receptors: simultaneous targeting of NMDARs and 5-HT_4_Rs exerts additive effects against stress

**DOI:** 10.1101/2023.09.27.559065

**Authors:** Briana K. Chen, Victor M. Luna, Michelle Jin, Abhishek Shah, Margaret E. Shannon, Michaela Pauers, Brenna L. Williams, Vananh Pham, Holly C. Hunsberger, Alain M. Gardier, Indira Mendez-David, Denis J. David, Christine A. Denny

## Abstract

**BACKGROUND:** Serotonin (5-HT) receptors and *N*-methyl-D-aspartate receptors (NMDARs) have both been implicated in the pathophysiology of depression and anxiety disorders. Here, we evaluated whether targeting both receptors through combined dosing of (*R*,*S*)-ketamine, an NMDAR antagonist, and prucalopride, a serotonin type IV receptor (5-HT_4_R) agonist, would have additive effects, resulting in reductions in stress-induced fear, behavioral despair, and hyponeophagia.

**METHODS:** A single injection of saline (Sal), (*R*,*S*)-ketamine (K), prucalopride (P), or a combined dose of (*R*,*S*)-ketamine and prucalopride (K+P) was administered before or after contextual fear conditioning (CFC) stress in both sexes. Drug efficacy was assayed using the forced swim test (FST), elevated plus maze (EPM), open field (OF), marble burying (MB), and novelty-suppressed feeding (NSF). Patch clamp electrophysiology was used to measure the effects of combined drug on neural activity in hippocampal CA3. c-fos and parvalbumin (PV) expression in the hippocampus (HPC) and medial prefrontal cortex (mPFC) was examined using immunohistochemistry and network analysis.

**RESULTS:** We found that a combination of K+P, given before or after stress, exerted additive effects, compared to either drug alone, in reducing a variety of stress-induced behaviors in both sexes. Combined K+P administration significantly altered c-fos and PV expression and network activity in the HPC and mPFC.

**CONCLUSIONS:** Our results indicate that combined K+P has additive benefits for combating stress-induced pathophysiology, both at the behavioral and neural level. Our findings provide preliminary evidence that future clinical studies using this combined treatment strategy may prove advantageous in protecting against a broader range of stress-induced psychiatric disorders.

## INTRODUCTION

Affective disorders are among the leading causes of global disease burden and have significantly increased in prevalence since the start of the COVID-19 pandemic (1). It is estimated that in 2020 alone, there was a 27.6% global increase in major depressive disorder (MDD) as well as an additional 25.6% surge in anxiety disorders. These rising numbers underscore the need for more effective prevention and treatment options for stress-related psychiatric disorders.

Current first-line treatments for anxiety and depression, such as selective serotonin reuptake inhibitors (SSRIs), largely modulate serotonergic signaling in the brain. Discovered several decades ago, the efficacy of SSRIs in reducing depressive symptoms significantly contributed to the monoamine hypothesis of mood disorders (2). This hypothesis suggests that antidepressants exert their efficacy by increasing the extracellular availability of monoaminergic neurotransmitters, including serotonin. Although SSRIs broadly increase serotonergic tone, emerging evidence suggests that specifically activating serotonin type IV receptors (5-HT_4_Rs) can provide more immediate, efficacious therapeutic benefits (3). In preclinical models, 5-HT_4_R agonists rapidly suppress behavioral despair, anhedonia, and anxiety-like behavior (4–6). Moreover, 5-HT_4_R agonists have been shown to enhance resilience to stress (7). Despite these promising data, to date, SSRIs such as fluoxetine and sertraline remain the first-line treatment strategy for depression and anxiety. Nonetheless, despite the ubiquity of SSRIs, issues such as nonspecific side effects, and lack of efficacy in patients suffering from treatment-resistant depression (TRD) have led many researchers to search for alternative theories to the monoamine hypothesis of mood disorders.

Alternatively, the glutamatergic hypothesis of depression has emerged as a leading hypothesis for affective disorders. This theory proposes that depressive and anxiogenic states arise from an imbalance of excitatory and inhibitory neurotransmission, potentially due to the abnormal activity of *N*-methyl-D-aspartate receptors (NMDARs) or α-amino-3-hydroxy-5-methyl-4-isoxazolepropionic acid receptors (AMPARs) (8–10). The glutamatergic hypothesis is strongly supported by the discovery that (*R,S*)-ketamine (K), an NMDAR antagonist, exerts rapid-acting antidepressant effects in as little as two hours, relieves symptoms of depression for up to three weeks, and is effective in treating TRD (11–14). Esketamine, a stereospecific enantiomer of K, recently became the first novel antidepressant approved by the FDA in nearly 20 years (15–19). Our lab and others have shown that K can be administered prior to stress to prevent the onset of stress-induced behavioral despair and decrease learned fear (20–25). Moreover, we have also demonstrated that fluoroethylnormemantine (FENM), a novel NMDAR antagonist, can also be effective when administered prior to or after stress (26,27). Together, these data strongly suggest that targeting NDMARs can be an effective treatment strategy for stress-related psychiatric disorders. However, although NMDAR antagonists exert demonstrated antidepressant effects, their utility in reducing anxiety-like behavior is limited (28).

Here, we sought to determine whether simultaneously targeting NMDARs and 5-HT_4_Rs could be effective in suppressing a wide variety of stress-induced fear, behavioral despair, and anxiety-like behaviors. A single injection of Sal, K, prucalopride (P), or combined K+P at different dosage combinations was administered prior to or after CFC stress. A single combinatorial dose of K+P, given before or after stress, exerted additive effects in reducing a variety of stress-induced behaviors in both sexes. Combined K+P administration attenuated bursts of AMPAR-mediated excitatory post-synaptic currents (EPSCs) in hippocampal CA3 and selectively altered correlated c-fos and parvalbumin (PV) network activity in the medial prefrontal cortex (mPFC) and hippocampus (HPC). Together, these results indicate that combined K+P exerts additive behavioral and neural effects compared to administration of either drug alone, suggesting that combinatorial pharmacological treatments to simultaneously target NMDARs and 5-HT_4_R may provide additional anxiolytic effects for further preclinical and clinical study.

## METHODS AND MATERIALS

For a full description of Methods and Materials, please refer to the **Supplemental Methods** in **Supplement 1**.

### Drugs

A single injection of saline (0.9% NaCl), (*R,S*)-ketamine (Ketaset, Zoetis, Parsippany-Troy Hills, NJ), prucalopride (SML1371, Sigma-Aldrich, St. Louis, MO), or combined (*R,S*)-ketamine + prucalopride was administered once during the course of each experiment at approximately eight weeks of age. All drugs were prepared in physiological saline and administered intraperitoneally (i.p.) in volumes of 0.1 cc per 10 mg body weight.

## RESULTS

### Prophylactic (*R*,*S*)-ketamine + prucalopride exerts additive anxiolytic effects in male and female mice

We previously reported that K, an NMDAR antagonist, and P, a 5-HT_4_R agonist, are effective prophylactics against stress (7,23,27). Although both compounds attenuate learned fear and reduce behavioral despair, neither drug has previously been shown to affect perseverative behavior, exploratory behavior, or hyponeophagia. Here, we hypothesized that combined K+P administration may result in additive behavioral effects. Male mice were injected with Sal, K, P, or combined K+P. One week later, mice were administered 3-shock CFC followed by behavioral testing (**Figure 1A**).

**Figure 1.**
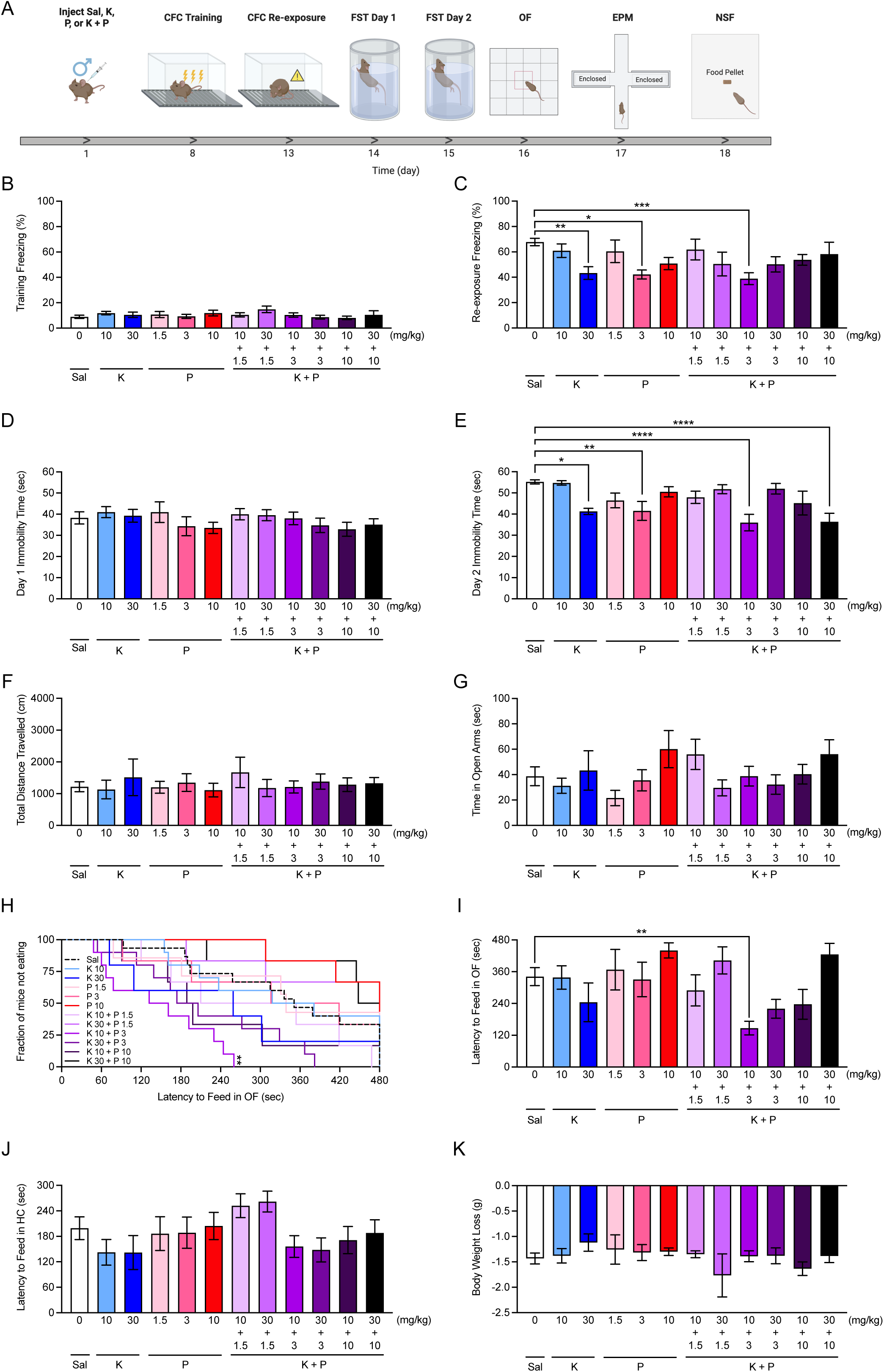
Combined prophylactic (*R,S*)-ketamine + prucalopride administration protects against stress in male 129S6/SvEv mice. **(A)** Experimental design. Sal, K, P, or combined K+P at different dosage combinations was administered one week prior to CFC stress in male 129S6/SvEv mice. **(B)** All groups of mice exhibited comparable freezing during CFC training. **(C)** Upon context re-exposure, mice administered K (30 mg/kg), P (3 mg/kg), or K+P (10 + 3 mg/kg) exhibited reduced freezing compared to Sal mice. **(D)** During day 1 of the FST day 1, all groups of mice exhibited comparable immobility time. **(E)** During day 2 of the FST, mice administered K (30 mg/kg), P (3 mg/kg), and K+P (10 + 3 or 30 + 10 mg/kg) exhibited decreased immobility time when compared to mice administered Sal. **(F)** All groups of mice traveled a comparable distance in the OF. **(G)** All groups of mice spent a comparable amount of time in the open arms of the EPM. **(H-I)** In the NSF, mice administered K+P (10 + 3 mg/kg) exhibited decreased latency to feed in the OF arena. **(J)** All groups of mice exhibited comparable latencies to feed in the HC. **(K)** All groups of mice loss a comparable amount of body weight following the NSF. (n = 5-15 male mice per group). Error bars represent ± SEM. * p < 0.05. ** p < 0.01. *** p < 0. 001, **** p < 0.0001. Sal, saline; K, (*R,S*)-ketamine; P, prucalopride; CFC, contextual fear conditioning; FST, forced swim test; OF, open field; EPM, elevated plus maze; NSF, novelty suppressed feeding; HC; home cage; cm, centimeter; min, minute; sec, second; mg, milligram; kg, kilogram.

All groups had comparable freezing during CFC training (**Figure 1B**). During re-exposure, K (30 mg/kg), P (3 mg/kg), or K+P (10 + 3 mg/kg), but no other doses, were effective at decreasing fear behavior when compared with Sal (**Figure 1C**). On day 2, but not on day 1 of the FST, K (30 mg/kg), P (3 mg/kg), or K+P (10 + 3 mg/kg and 30 + 10 mg/kg) decreased immobility time when compared with Sal (**Figure 1D-1E**).

Behavior was comparable across all groups in the OF and EPM (**Figure 1F-1G**) In the NSF, our prior work indicated that K and P separately were not effective at reducing hyponeophagia (7, 21, 25). However, combined K+P (10 + 3 mg/kg) decreased the latency to feed in the OF when compared with Sal without altering other measures of appetite or motivation (**Figure 1H-1K**). These data indicate that the combined K+P has a synergistic effect of decreasing stress-induced hyponeophagia in male mice with the additional effect of decreasing the dose of K needed to attenuate fear expression (i.e., 30 versus 10 mg/kg when combined with P).

To determine whether combined K+P, could have additive prophylactic effects in females, we administered the same injection and behavioral schedule to female mice as outlined in **Figure 1**, with the exception that the dose of K and P was chosen based on prior studies in female 129S6/SvEv mice (**Figure 2A**) (7,23). All groups had comparable freezing during CFC training and re-exposure (**Figure 2B**-**2C**). On day 1 of the FST, K (10 mg/kg), P (3 mg/kg), or combined K+P (10 + 1.5 mg/kg) reduced immobility time (**Figure 2D**). On day 2 of the FST, K (10 mg/kg), P (1.5 and 3 mg/kg), or combined K+P (10 + 1.5 mg/kg) reduced immobility time (**Figure 2E**). Behavior was comparable across all groups in the OF (**Figure 2F**). In the EPM, P (3 mg/kg) increased time in the open arms (**Figure 2G**). In the NSF, our prior work indicated that the 5HT_4_R agonist RS-67,333 was effective at reducing hyponeophagia in female 129S6/SvEv mice, but we have not yet previously tested if P was also effective at reducing hyponeophagia in female 129S6/SvEv mice. Here, only combined K+P (10 + 1.5 mg/kg) reduced the latency to feed in the OF without altering body weight loss or behavior in the home cage (**Figure 2H**-**2K**). These data indicate that combined K+P has an additive effect in reducing stress-induced hyponeophagia in female mice.

**Figure 2.**
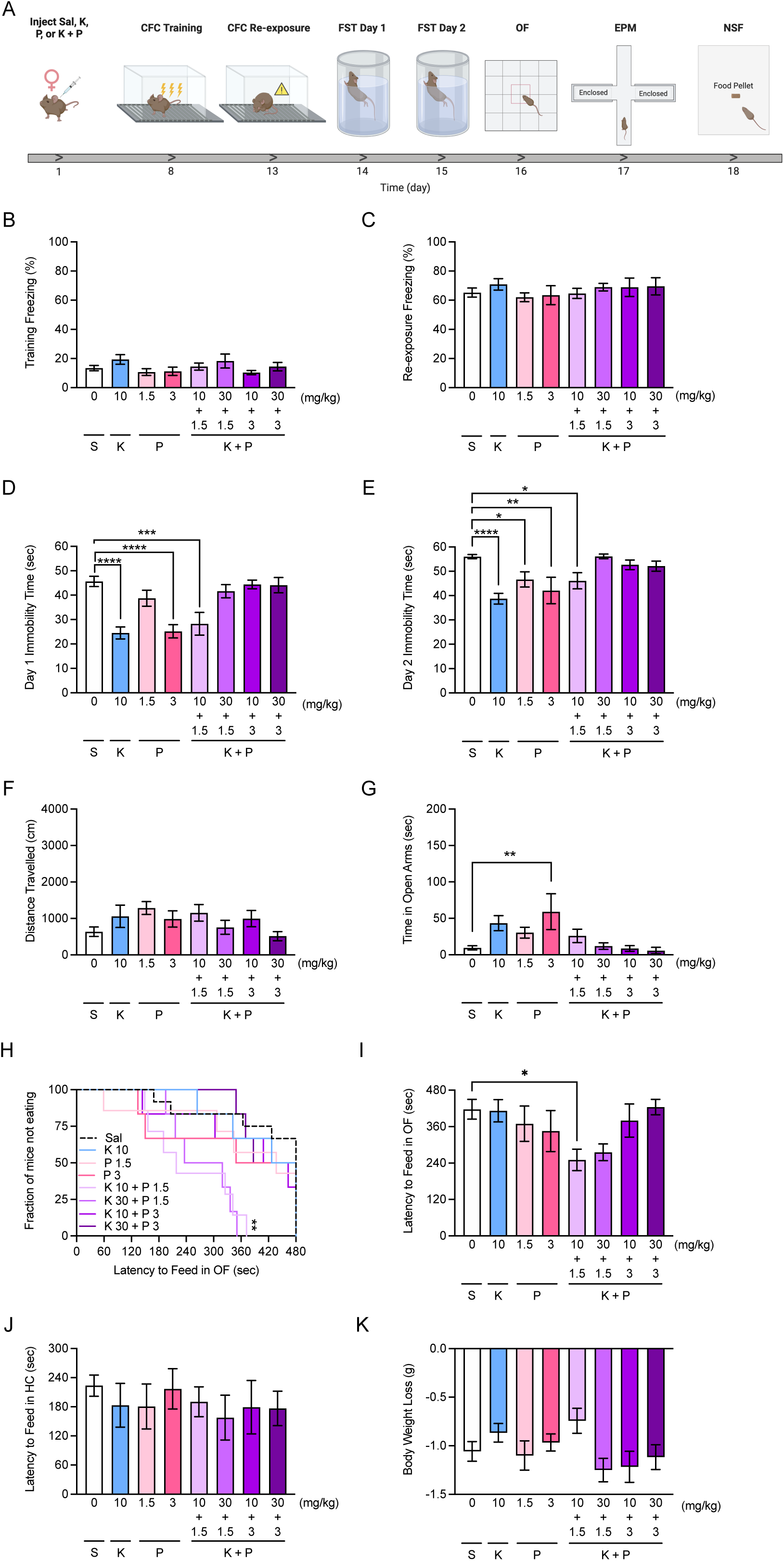
Combined prophylactic (*R,S*)-ketamine + prucalopride administration protects against stress in female 129S6/SvEv mice. **(A)** Experimental design. (B-C) All groups of mice exhibited comparable freezing during CFC training and re-exposure. **(D)** During day 1 of the FST, mice administered K (10 mg/kg), P (3 mg/kg), and K+P (10 + 1.5 mg/kg) exhibited decreased immobility time when compared to mice administered Sal. **(E)** During day 2 of the FST, mice administered K (10 mg/kg), P (3 or 10 mg/kg), and K+P (10 + 1.5 mg/kg) exhibited decreased immobility time when compared to mice administered Sal. **(F)** All groups of mice traveled a comparable distance in the OF. **(G)** P (3 mg/kg), but no other drug tested, significantly increased time spent in the open arms of the EPM when compared to Sal. **(H-I)** In the NSF, mice K+P (10 + 1.5 mg/kg) exhibited decreased latency to feed in the OF arena. **(J)** All groups of mice exhibited comparable latencies to feed in the HC. **(K)** All groups of mice loss a comparable amount of body weight following the NSF. (n = 6-12 female mice per group). Error bars represent ± SEM. * p < 0.05. ** p < 0.01. *** p < 0. 001, **** p < 0.0001. Sal, saline; K, (*R,S*)-ketamine; P, prucalopride; CFC, contextual fear conditioning; FST, forced swim test; OF, open field; EPM, elevated plus maze; NSF, novelty suppressed feeding; HC; home cage; cm, centimeter; min, minute; sec, second; mg, milligram; kg, kilogram.

### Prophylactic (*R,S*)-ketamine + prucalopride facilitates contextual fear discrimination in male, but not female mice

We previously demonstrated that prophylactic administration of K facilitates and enhances CFD in in male mice (29). However, it is still unknown whether our combined K+P findings extend to other models of stress. Here, we administered Sal, K, P, or K+P one week prior to a CFD paradigm in male and female 129S6/SvEv mice (**Figure S1A**).

In male mice, combined K+P accelerated CFD (**Figure S1B-S1I**). Specifically, only mice administered combined K+P could discriminate between the contexts on day 4 of the CFD paradigm (**Figure S1G**). In female mice, K and P both accelerated CFD, but not when administered as combined K+P (**Figure S1J-S1Q**). All mice discriminated by day 10 of the CFD paradigm. In summary, these data suggest that combined K+P administration can slightly facilitate fear discrimination in male, but not female mice.

### (*R*,*S*)-ketamine + prucalopride reduces perseverative behavior and hyponeophagia in non-stressed female, but not male mice

To determine whether combined K+P alters behavior in non-stressed mice, we administered Sal, K, P, or K+P and then administered the FST 1 hour later (**Figure S2A**). In male mice, immobility time was not attenuated by drug administration (**Figure S2B-S2E**). In female mice, overall, but not average, immobility time on both Days 1 and 2 was increased by K+P administration (**Figure S2F-S2I**).

Next, we tested whether combined K+P alters exploratory, perseverative, or hyponeophagia behaviors in non-stressed mice. Sal, K, P, or K+P was administered one hour prior to the OF. The EPM, MB, and NSF assays were administered on subsequent days (**Figure S3A**). In male mice, behavior in the OF, EPM, MB, and NSF was not significantly altered by drug administration (**Figure S3B-S3I**). In female mice, P significantly increased locomotion in the OF when compared to Sal (**Figure S3J-S3K**). Behavior in the EPM was comparable across all drug groups (**Figure S3L-S3M**). In the MB assay, mice administered K+P buried significantly fewer marbles compared to mice administered Sal (**Figure S3N**). In the NSF assay, combined K+P reduced the latency to feed in the OF without altering the latency to feed in the home cage (**Figure S3O-S3Q**). These results indicate that combined K+P reduces perseverative behavior and hyponeophagia in non-stressed female, but not male mice.

### Prophylactic (*R*,*S*)-ketamine + prucalopride attenuates AMPAR-mediated bursts of excitatory activity in hippocampal CA3

We previously demonstrated that prophylactic drugs, including K and P, diminish large-amplitude AMPAR-mediated excitatory post-synaptic currents (EPSCs) (7,23,27). Here, we hypothesized that combined K+P would also inhibit AMPAR bursting. Sal or combined K+P was administered and one week later, mice were sacrificed, and patch clamp electrophysiology was performed in hippocampal CA3 (**Figure 3A**). While Sal-administered mice exhibited bursts of large-amplitude AMPAR-mediated EPSCs (**Figure 3B-3C**), K+P-administered mice did not exhibit these AMPAR bursts in CA3 (**Figure 3E-3F**). Combined K+P administration led to a decrease in mean EPSC amplitude (**Figure 3D**) as well as a trending, but not significant reduction in the number of EPSCs (**Figure 3G**). Together, these data show that, similarly to administration of K or P alone, combined K+P also inhibits large-amplitude bursts of AMPAR-mediated excitatory activity in hippocampal CA3.

**Figure 3.**
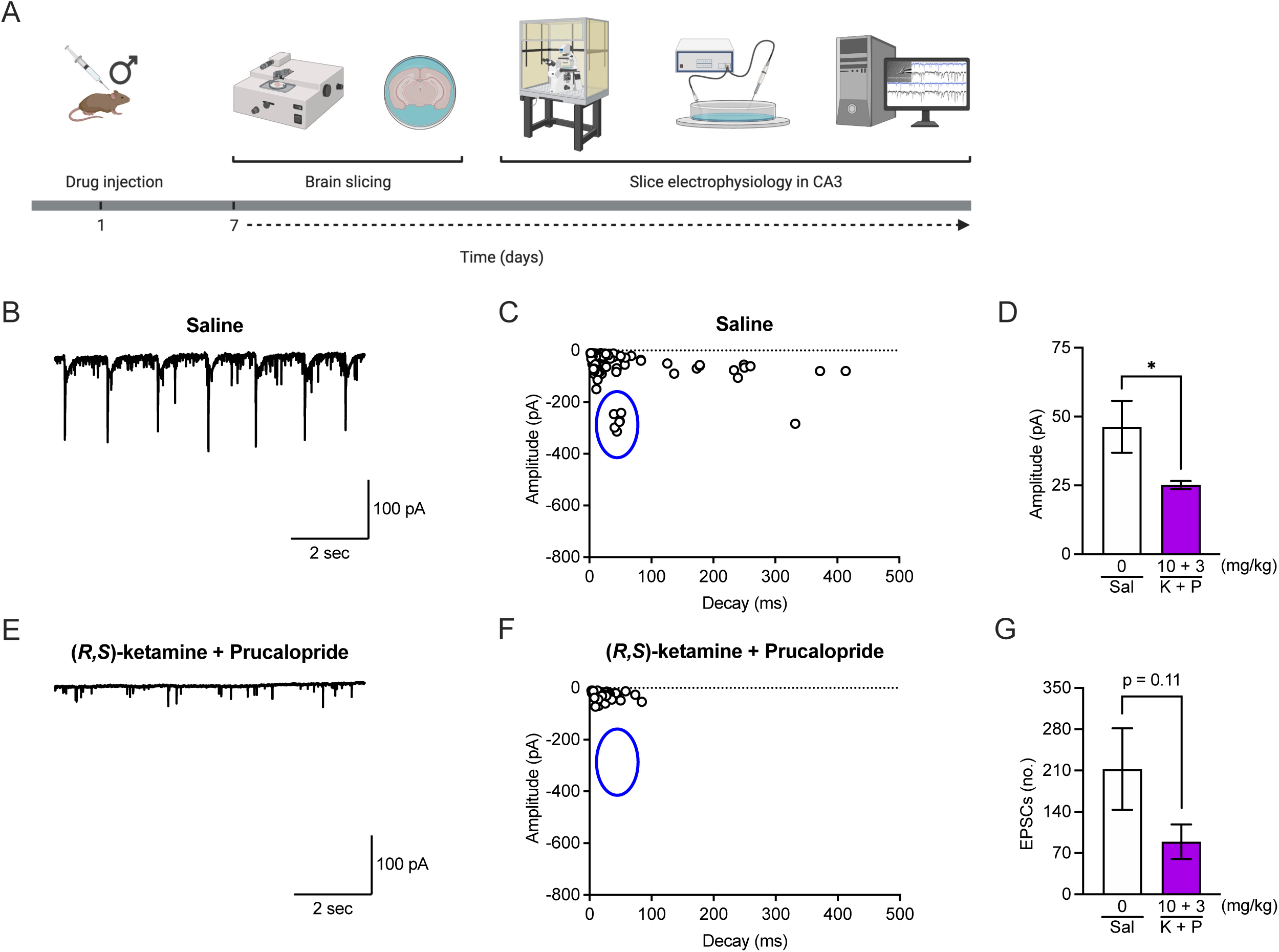
Combined (*R,S*)-ketamine + prucalopride blocks reduces AMPAR-mediated bursting in hippocampal CA3. **(A)** Experimental design. Male 129S6/SvEv mice were i.p. injected with Sal or K+P (10 + 3 mg/kg) and sacrificed for electrophysiology one week later. Representative EPSCs in **(B)** Sal-and **(E)** K+P-administered mice. **(C)** Sal-administered mice exhibited bursts of large-amplitude AMPAR-mediated activity (circled in blue) which were blocked in **(F)** K+P-administered mice. **(D)** Mean EPSC amplitude was significantly reduced by K+P administration. **(G)** There was also a trending, but not significant, decrease in number of EPSCs in mice given K+P when compared with Sal. (n = 5-7 cells per group). Error bars represent ± SEM. * p < 0.05. Sal, saline; K+P, (*R,S*)-ketamine + prucalopride; CA3, Cornu ammonis 3; sec, second; pA, picoampere; mg, milligram; kg, kilogram; ms, millisecond; EPSC, excitatory post-synaptic current.

### Prophylactic (*R*,*S*)-ketamine + prucalopride selectively reduces excitatory signaling in mPFC and HPC

Our lab has previously reported that prophylactic K administration upregulates c-fos expression in vCA3 during CFC re-exposure (29). Here, we hypothesized that combinatorial K+P administration could alter neural signaling along the dorsoventral axis of the HPC as well as in the mPFC, a functionally connected downstream brain region that, along with the HPC, regulates memory encoding, memory retrieval, emotional processing, and the stress response (30–32). Male mice were administered Sal, K, P, or K+P and 1 week later, administered CFC and the FST. One hour later, mice were sacrificed, and brains were processed for c-fos immunoreactivity (**Figure 4A-4B**). As previously demonstrated, K, P, and K+P significantly attenuated learned fear and behavioral despair in male mice **(Figure S4A-S4D**).

**Figure 4.**
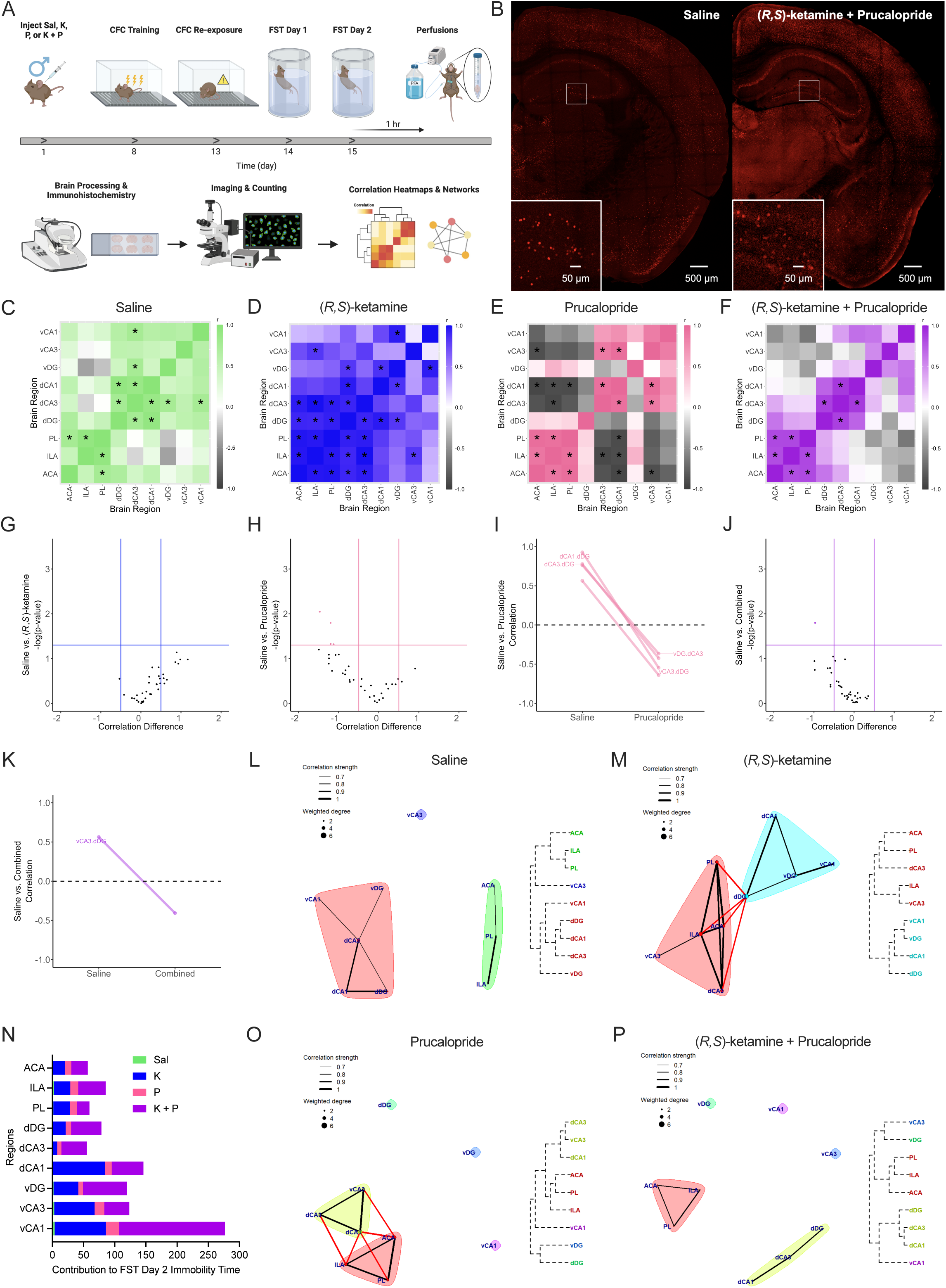
Combined (*R,S*)-ketamine + prucalopride alters correlated excitatory signaling in the mPFC and HPC. **(A)** Behavioral paradigm. Mice were administered a single prophylactic injection of Sal, K, P, or combined K+P one week prior to CFC stress. Five days later, mice were re-exposed to the training context and tested in the FST. Mice were sacrificed one hour after day 2 of the FST, and immunohistochemistry was used to quantify c-fos and PV expression. **(B)** Representative images of c-fos immunostaining in (left) Sal-and (right) combined K+P-administered mice. Insets reveal close-ups of hippocampal c-fos expression. Heat map correlations of c-fos expression in **(C)** Sal-, **(D)** K-, **(E)** P-, and **(F)** combined K+P-administered mice. Green, blue, pink, or purple indicate strong positive correlations, while gray indicates strong negative correlations. **(G)** Volcano plot indicating regional correlation differences greater than 0.5 between Sal-and K-administered mice. **(H-I)** Volcano and parallel plots indicating regional correlation differences greater than 0.5 between Sal-and P-administered mice. **(J-K)** Volcano and parallel plots indicating regional correlation differences greater than 0.5 between Sal-and combined K+P-administered mice. Notably, mice given combined K+P exhibited a decrease in correlated vCA3-dDG activity when compared with Sal. Cluster maps of correlated c-fos expression in **(L)** Sal-, **(M)** K-, **(O)** P-, and **(P)** combined K+P-administered mice. The cluster map reveals strongly interconnected activity in Sal-, K-, and P-administered mice. In contrast, the combined K+P-administered cluster map is less cohesive, with the most isolation in all ventral hippocampal regions. **(N)** A bootstrap prediction analysis revealed that vCA1 and vDG contribute the most to immobility time during FST Day 2 in K+P-administered mice. (n = 7-8 mice per group). Error bars represent ± SEM. * p < 0.05. Sal, saline; K, (*R,S*)-ketamine; P, prucalopride; K + P, (*R,S*)-ketamine + prucalopride; CFC, contextual fear conditioning; FST, forced swim test; μm, micrometers; ACA, anterior cingulate area; ILA, infralimbic area; PL, prelimbic area; dDG, dorsal dentate gyrus; dCA3, dorsal field CA3; dCA1, dorsal field CA1; vDG, ventral dentate gyrus; vCA3, ventral field CA3; vCA1, ventral field CA1.

Heatmap correlations between selected brain regions revealed an increase in negative correlations in P-and combined K+P-administered mice when compared with saline mice (**Figure 4C-4F**). Using volcano and parallel plots, we confirmed this increase in negative correlations (**Figure 4G-4K, S4E-S4J**). Notably, vCA3 to dDG was the only significantly negative correlation in mice given combined K+P (**Figure 4K**).

We then aimed to determine the functional consequences of altered c-fos expression in the mPFC and HPC. First, we generated cluster network maps to visualize functionally connected subregions of the HPC and mPFC during FST Day 2 (**Figure 4L-4M, 4O-4P**). In mice administered combined K+P, vHPC was disconnected from other regions, but there was interconnectivity within mPFC and within dHPC. This isolation of the vHPC was not observed in Sal mice or in K-or P-administered mice. Next, using a bootstrap prediction analysis, we aimed to identify the regions contributing most to immobility during FST Day 2 (**Figure 4N**). In Sal-administered mice, vCA3 and vCA1 contributed most to immobility time. dCA1 and vCA3 contributed most in K-administered mice, while vCA3 and ACA contributed most to immobility in P-administered mice. In combined K+P-administered mice, vCA1 and vDG gave the greatest contributions to immobility time. These data establish vHPC as a critical modulator of stress-related behavior in the FST.

Next, we modeled the data as a network, with regions acting as nodes and correlations functioning as edges, then used network analysis to assess circuit-level connectivity during day 2 of the FST (**Figure 5A-5D**). Combined K+P resulted in a sparser network when compared to the three other groups (**Figure 5D**), suggesting a refinement in functionally connected excitatory activity in mice administered combined drug. Interestingly, in experimental drug groups, there was an enhancement of interconnectivity within mPFC subregions, particularly between ACA and ILA. K, but not any other drug, significantly increased mean degree, global efficiency, and distance of the network when compared with control saline (**Figure 5E-5F, 5I**). Mean clustering coefficient and betweenness centrality was comparable across all drug groups (**Figure 5G-5H**). Interestingly, combined K+P, but not K or P alone, significantly increased c-fos expression across the mPFC as well as in dCA3, dCA1, and vCA3 (**Figure 5J-5O**). These results suggest that combined K+P significantly increases excitatory activity in mPFC and HPC.

**Figure 5.**
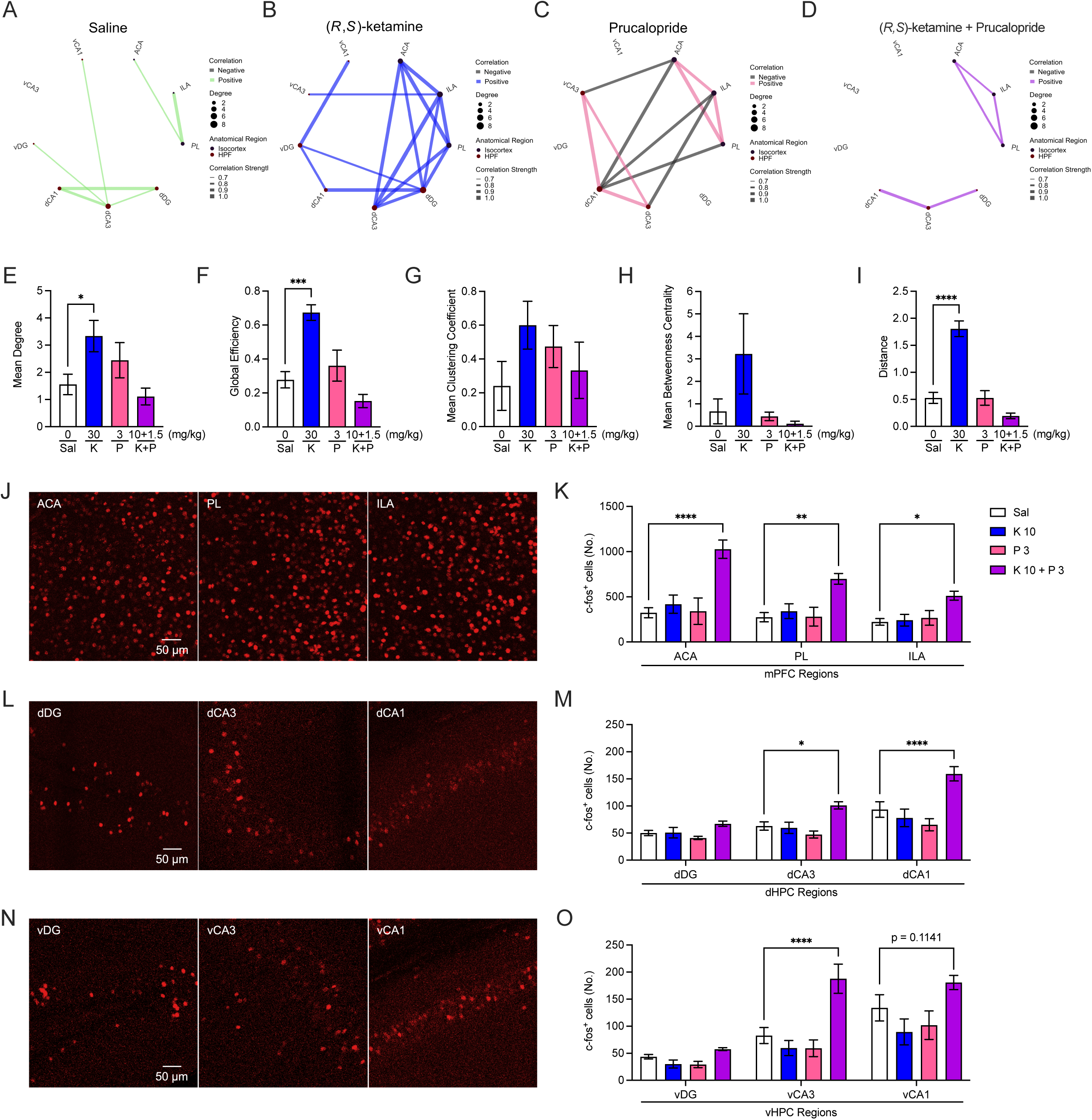
Combined (*R,S*)-ketamine + prucalopride alters network c-fos activity in the mPFC and HPC. c-fos expression modeled as networks of functional activity in **(A)** Sal-, **(B)** K-, **(C)** P-, and **(D)** combined K+P-administered mice. Green, blue, pink, or purple indicate positive correlations, while gray indicates strong negative correlations. Thicker lines indicate stronger correlations and larger circles represent an increased degree of nodes. Combined K+P administration led to sparser network activity when compared with Sal, K, and P administration. **(E-F)** Mean degree and global efficiency are significantly increased in K-administered, but not P-or combined K+P-administered mice, when compared to Sal-administered mice. **(G-H)** Mean clustering coefficient and betweenness centrality are comparable across all drug groups. **(I)** K significantly increases network distance in comparison to Sal. Representative images of c-fos immunostaining in **(J)** ACA, PL, and ILA of the mPFC, **(L)** dDG, dCA3, and dCA1 of the dHPC, and **(N)** vDG, vCA3, and vCA1 of the vHPC. When compared with Sal, (*K+P* significantly increased the number of c-fos^+^ cells in **(K)** mPFC, **(M)** dCA3, dCA1, and **(O)** vCA3. (n = 7-8 mice per group). Error bars represent ± SEM. * p < 0.05, ** p < 0.01, *** p < 0. 001, **** p < 0.0001. ACA, anterior cingulate area; ILA, infralimbic area; PL, prelimbic area; dDG, dorsal dentate gyrus; dCA3, dorsal field CA3; dCA1, dorsal field CA1; vDG, ventral dentate gyrus; vCA3, ventral field CA3; vCA1, ventral field CA1; Sal, saline; K, (*R,S*)-ketamine; P, prucalopride; K + P, (*R,S*)-ketamine + prucalopride; mg, milligram; kg, kilogram; μm, micrometers; no., number; mPFC, medial prefrontal cortex; dHPC, dorsal hippocampus; vHPC, ventral hippocampus.

### Prophylactic (*R*,*S*)-ketamine + prucalopride selectively enhances inhibitory signaling in mPFC and HPC

To better understand how inhibitory signaling is altered by drug administration, we examined expression of PV, a calcium-binding protein that is expressed in fast-spiking inhibitory interneurons, because it plays a critical role in modulating neural activity, and because its expression is sensitive to stress and antidepressant administration (33–35). First, we examined heatmap correlations of PV expression in the mPFC, dHPC, and vHPC followed by volcano and parallel plots (**Figure 6A-6D, S5A-S5K**). Combined K+P significantly increased positively correlated PV expression (**Figure 6D**), particularly between vCA1 and vDG as well as vDG and dCA1 (**Figure S5E**). We then used cluster network maps to reveal functionally correlated PV expression across brain regions (**Figure S5L-S5O**). Combined K+P, in comparison to Sal, enhanced functional connectivity between a majority of dHPC and vHPC regions (**Figure S5L, S5O**). Interestingly, ILA and vCA3 were functionally isolated from other areas of the brain.

Subsequently, we modeled correlated PV expression as an inhibitory network, with the regions as nodes, and correlated expression as edges (**Figure 6E-6H**). Mice administered combined K+P showed an increase in correlated inhibitory expression across the HPC when compared with Sal; in particular, mPFC regions were isolated from both dorsal and ventral HPC regions. Computed network measures showed that mean degree was comparable across all groups (**Figure S5P**). Global efficiency, mean betweenness centrality, and distance measures were significantly increased in P-administered mice, and mean clustering coefficient was significantly reduced in K-administered mice (**Figure S5Q-S5T**). Interestingly, when looking at PV expression across the mPFC and the HPC, combined K+P selectively increased the number of PV cells in vCA3, but not in any other subregion (**Figure 6L-6N**). These data suggest that although combined K+P may enhance PV expression in a select subregion of the vHPC, it may still enhance correlated inhibitory tone throughout the mPFC and HPC.

**Figure 6.**
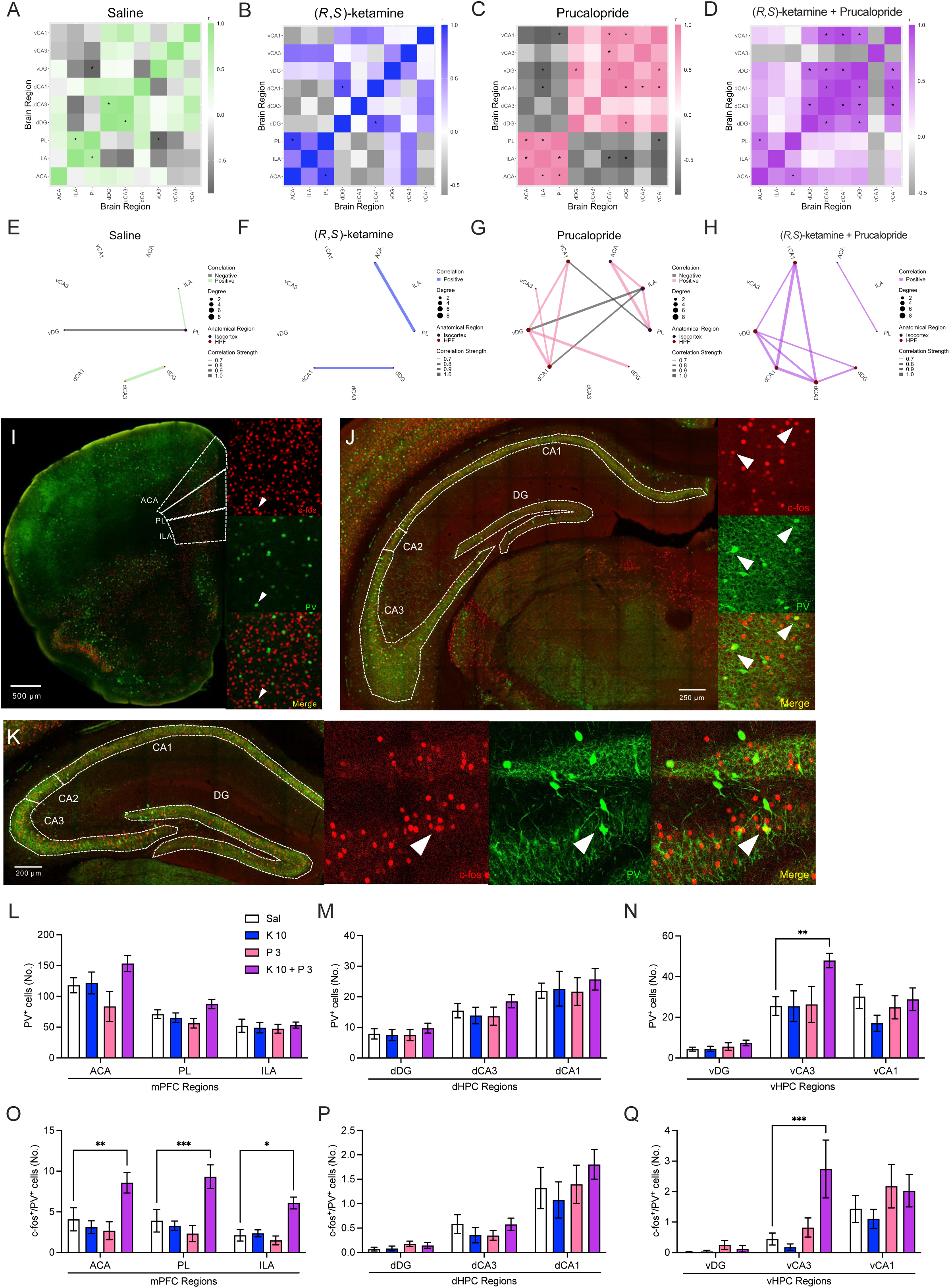
Combined (*R,S*)-ketamine + prucalopride selectively enhances correlated PV expression. Heat map correlations of PV expression in **(A)** Sal-, **(B)** K-, **(C)** P-, and **(D)** combined K+P-administered mice. Green, blue, pink, or purple indicate strong positive correlations, while gray indicates strong negative correlations. PV expression modeled as networks of functional activity in **(E)** Sal-, **(F)** K-, **(G)** P-, and **(H)** combined K+P-administered mice. Green, blue, pink, or purple indicate positive correlations, while gray indicates strong negative correlations. Thicker lines indicate stronger correlations and larger circles represent an increased degree of nodes. Combined K+P administration led to increased inhibitory network connectivity when compared with Sal administration. Representative images of c-fos and PV immunostaining in **(I)** mPFC, **(J)** vHPC, and **(K)** dHPC. The number of PV^+^ cells is comparable in all subregions of the **(L)** mPFC and **(M)** dHPC. **(N)** Combined K+P increases PV expression in vCA3, but no other subregion of the vHPC. **(O)** Combined drug administration significantly increased the number of c-fos^+^/PV^+^ co-labeled cells across the mPFC. **(P)** The number of c-fos^+^/PV^+^ co-labeled cells was comparable across the dHPC in all groups. **(Q)** Combined K+P increased the number of PV^+^/c-fos^+^ co-labeled cells in vCA3, but no other subregion of the vHPC. (n = 7-8 mice per group). Error bars represent ± SEM. * p < 0.05, ** p < 0.01, *** p < 0. 001. ACA, anterior cingulate area; ILA, infralimbic area; PL, prelimbic area; dDG, dorsal dentate gyrus; dCA3, dorsal field CA3; dCA1, dorsal field CA1; vDG, ventral dentate gyrus; vCA3, ventral field CA3; vCA1, ventral field CA1. μm, micrometers; mPFC, medial prefrontal cortex; dHPC, dorsal hippocampus; vHPC, ventral hippocampus.

### Prophylactic (*R*,*S*)-ketamine + prucalopride increases c-fos and PV co-localization in the mPFC and vCA3

Finally, we examined co-expression of c-fos and PV in the mPFC and HPC (**Figure 6I-6K**). Across all subregions of the mPFC, combined K+P increased the number of PV^+^/c-fos^+^ cells when compared to Sal (**Figure 6O**). In the dHPC, all groups exhibited comparable levels of PV^+^/c-fos^+^ cells (**Figure 6P**). In the vHPC, only combined K+P significantly increased the number of PV^+^/c-fos^+^ cells in vCA3, but not in the vDG or vCA1 (**Figure 6Q**). These data suggest that simultaneous targeting of the NMDAR and 5-HT_4_R increases the activity of PV^+^ cells in the mPFC and vCA3.

### Combined (*R*,*S*)-ketamine + prucalopride is effective when administered after stress

Recently, we demonstrated that administration of an NMDAR antagonist after stress prevents stress-induced behavioral despair in male and female mice (27). To test whether simultaneously targeting NMDARs and 5HT_4_Rs could also be protective when administered following stress, we administered a single dose of Sal, K, P, or K+P five minutes after CFC (**Figure 7A**).

**Figure 7.**
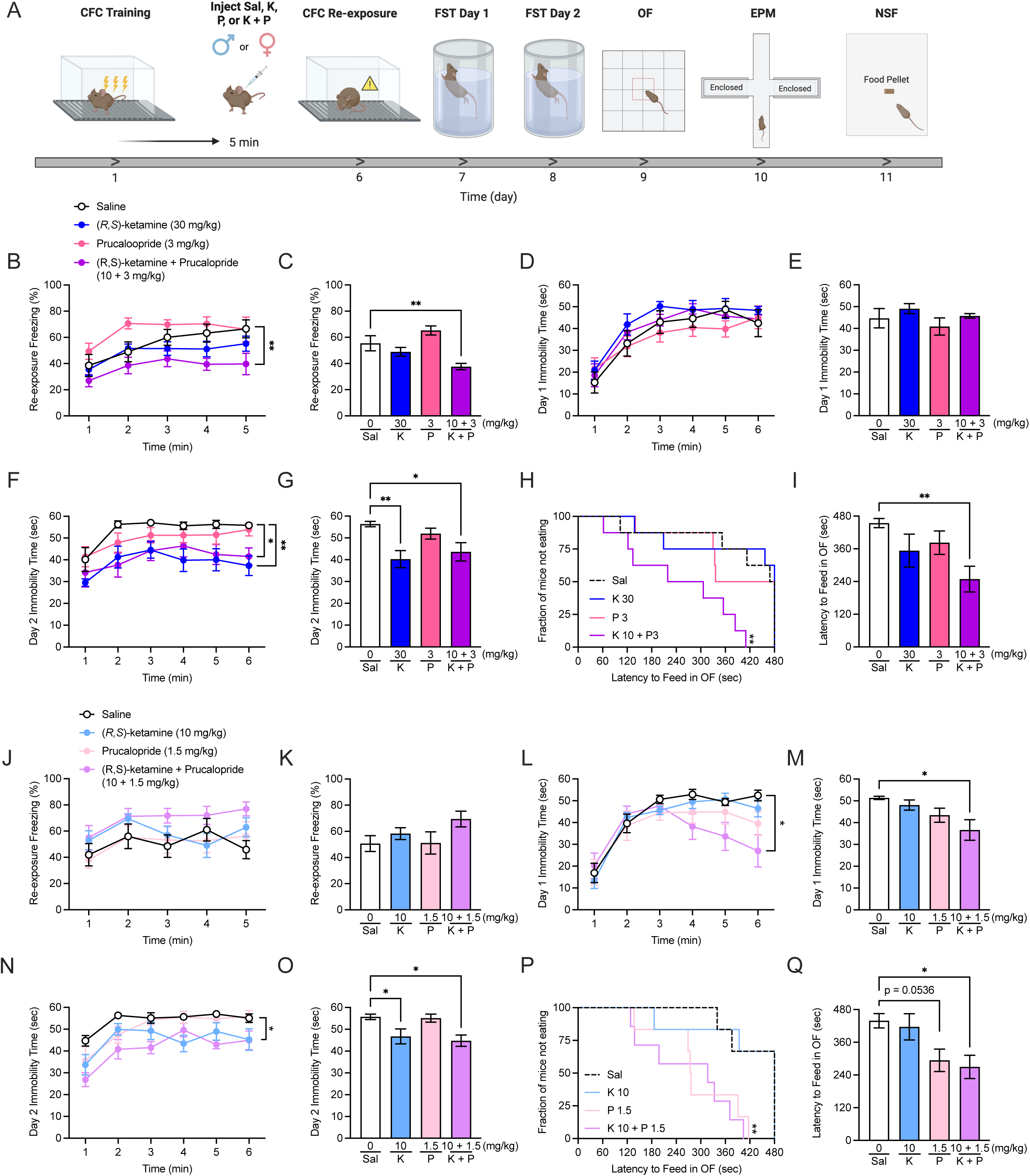
Combined (*R,S*)-ketamine + prucalopride attenuates learned fear in male mice and reduces behavioral despair and hyponeophagia in both sexes when administered after stress. **(A)** Experimental design. Male and female 129S6/SvEv mice were administered a single administration of Sal, K, P, or a combined K+P five minutes after 3-shock CFC. **(B-C)** In male mice, during context re-exposure, combined K+P, but not K or P alone, significantly reduced freezing when compared to Sal. **(D-E)** During day 1 of the FST, all groups of mice exhibited comparable immobility. **(F-G)** During day 2 of the FST, male mice administered K or combined K+P exhibited decreased immobility time when compared to Sal. **(H-I)** In the NSF, mice administered combined K+P exhibited a reduced latency to feed in the OF when compared with mice administered Sal. **(J-K)** In female mice, during context re-exposure, freezing was comparable across all groups. **(L-M)** During day 1 of the FST, mice administered combined K+P exhibited decreased immobility time when compared to Sal. **(N-O)** During day 2 of the FST, female mice administered K or K+P exhibited decreased immobility time when compared to Sal. **(P-Q)** In the NSF, mice administered combined K+P exhibited decreased latency to feed when compared to Sal. (n = 5-7 male or female mice per group). Error bars represent ± SEM. * p < 0.05. ** p < 0.01. *** p < 0. 001, **** p < 0.0001. CFC, contextual fear conditioning; FST, forced swim test; OF, open field; EPM, elevated plus maze; NSF, novelty suppressed feeding; mg, milligram; kg, kilogram; Sal, saline; K, (*R,S*)-ketamine; P, prucalopride; K + P, (*R,S*)-ketamine + prucalopride; min, minute; sec, second.

In male mice, combined K+P, but not single drug administration alone, significantly decreased learned fear (**Figure 7B-7C**), behavioral despair on day 2, but not day 1 of the FST (**Figure 7D-7G**), and hyponeophagia (**Figure 7H-7I**). Of note, combined K+P did not affect behavior in the OF, EPM, or other measures during the NSF (**Figure S6A-S6L**).

In female mice, combined K+P did not alter learned fear (**Figure 7J-7K**). On days 1 and 2 of the FST, combined K+P decreased behavioral despair (**Figure 7L-7O**). In the NSF, combined K+P decreased hyponeophagia (**Figure 7P-7Q**). Of note, combined K+P administration did not affect behavior in the OF, EPM, or other measures during the NSF (**Figure S6M-S6X**). Overall, these data indicate that combined K+P is effective in preventing a variety of stress-induced behaviors in male and female mice when administered after stress.

## DISCUSSION

Here, we characterized the behavioral and neural effects of combined K+P administration. We discovered that simultaneous targeting of NMDARs and 5-HT_4_Rs prior to or after stress exerts additive protective effects against stress-induced behaviors in male and female mice. Combined K+P, but not either drug administered alone, significantly enhanced c-fos and PV expression in the mPFC and select regions of the HPC. Overall, our results suggest that the simultaneous targeting of NMDARs and 5-HT_4_Rs using a K+P drug combination exerts additional and distinct neural and behavioral benefits compared to administration of a single drug.

To our knowledge, combinatorial targeting of NMDARs and 5-HT_4_Rs has not previously been studied; clinical studies have shown that adjunctive administration of Spravato^®^ in addition to continued SSRI treatment is an effective therapeutic strategy for patients suffering from TRD and suicidal ideation (15–18). Thus, implementing combined drug administration in the clinic is a tractable and applicable method of treatment for patients suffering from psychiatric disorders. Behaviorally, we found that combined K+P was effective in reducing hyponeophagia in both male and female mice. These results are critical, as they indicate that simultaneously targeting NMDARs and 5-HT_4_Rs can suppress a larger variety of anxiety-related phenotypes compared to targeting either receptor alone. In particular, the NSF assay is a measure of the degree to which stress (e.g., a novel environment) can affect feeding behavior and is purported to quantify hyponeophagia (36,37). Our data suggest that although K and P are not reported to affect clinical symptoms of anxiety, when combined, they may be an effective treatment option for further study (28,38).

We demonstrated a strong dose specificity of the K+P combination. Although we initially hypothesized that combining the behaviorally-effective doses of (*R,S*)-ketamine (30 mg/kg in male mice or 10 mg/kg in female mice) and prucalopride (3 mg/kg in male mice or 1.5 mg/kg in female mice) would exhibit the strongest behavioral effect, our experiments indicated that the most effective dose was 10 mg/kg of (*R,S*)-ketamine combined with 3 or 1.5 mg/kg of prucalopride in male or female mice, respectively. This unexpected result shows that there is a specific drug concentration sufficient to both block NMDARs and activate 5-HT_4_Rs to an optimal degree. Our data suggest that combined K+P may have an inverted U-shaped dose-effect curve that is common in many active compounds (39). Further experimentation may shed light on this phenomenon by testing a greater range of K+P doses and examining potential mechanisms contributing to the drug combination’s dose-specific effect.

Previously, we have demonstrated that a variety of prophylactic drugs, including NMDAR antagonists and 5-HT_4_R agonists, block bursts of large-amplitude AMPAR-mediated EPSCs in CA3 (7,23,27). Our results indicate that combined K+P administration also attenuates AMPAR-mediated bursts of activity in CA3. The spontaneous firing and large amplitude of these AMPAR-mediated bursts closely resemble the characteristic features of hippocampal sharp wave activity (SPW), which plays an important role in memory formation and sleep (40–43). Sharp wave ripples (SPW-Rs) emerge from the combined synchronous activity of a small subset of excitatory pyramidal cells and inhibitory interneurons, particularly PV^+^ basket cells, in CA3 (41,42,44). These hippocampal neural events are critical for memory encoding, consolidation, and retrieval and may function to link emotional salience with contextual information (45,46). SPW-Rs are reported to trigger long-lasting synaptic depression which may help to refine the specificity of memory engrams (47). K has been previously shown to reduce the occurrence of SPW-Rs in CA1 up to 30 minutes after administration (48,49). Our results suggest that this K-induced suppression of SPW-Rs may also occur in CA3 and may last for up to one week after administration. Furthermore, CA3 SPW-Rs are suppressed by high 5-HT levels (50). As 5-HT_4_R agonists may increase the release of 5-HT, P may also suppress hippocampal SPW-Rs by increasing serotonergic tone (51). Our data suggest that targeting both NMDARs and 5-HT_4_Rs together, along with targeting either receptor individually, may block hippocampal SPW-Rs in CA3 for up to 1 week. Functionally, this suppression of SPW-Rs may cognitively decouple the contextual experience of a stressor with negative valence, allowing for subsequent memory retrieval of the event without debilitating fear and preventing associated symptoms of affective disorders. However, further study is necessary to test the validity of this hypothesis.

The behavioral and electrophysiological results of our combined drug strategy are complemented by our network analysis of excitatory and inhibitory signaling in the mPFC and HPC. These findings revealed that although prophylactic K, P, or combined K+P result in similar behavioral consequences, they may exert distinct effects on correlated c-fos and PV expression. Curiously, K+P appears to exert similar but still distinct effects on c-fos and PV network activity as P. Notably, K+P reduces correlated c-fos network activity, but only between vCA3 and DG, while increasing c-fos expression in the mPFC, select regions of the dHPC, and vCA3. Combined K+P also increased correlated PV expression, but to a lesser extent than P alone, and upregulated PV^+^ neurons selectively in vCA3. These results suggest that simultaneous NMDAR and 5-HT_4_R targeting can refine excitatory/inhibitory (E/I) balance in vCA3, which may contribute to the suppression of SPW-Rs and subsequently affect neural activity in downstream brain regions, thus improving resilience to stress.

In conclusion, we report that combined K+P exerts synergistic effects in reducing stress-induced fear, behavioral despair, and hyponeophagia behaviors in both male and female mouse models of stress. Simultaneously targeting NMDARs and 5-HT_4_Rs is sufficient to modulate both excitatory and inhibitory signaling in the mPFC and HPC, brain regions critically involved in stress processing. Nonetheless, further study is necessary to elucidate the molecular mechanisms and clinical efficacy of simultaneous NMDAR antagonist and 5-HT_4_R agonist administration. Overall, the present study demonstrates the potential of utilizing adjunctive pharmacological treatment to advance targeted therapies for stress-related psychiatric disorders.

## ACKNOWLEDGEMENTS AND DISCLOSURES

BKC was supported by an NIMH F31 MH121023. MJ was supported by the MD-PhD program at Columbia University and a T32 GM7367-46. AS and MP were supported by an NIMH T32 MH1226036. BLW was supported by an NHLBI T32 HL120826. VP was supported by an NICHD R01 HD101402. HCH was supported by an NIA K99 AG059953 and an NIA R24 AG061421. AMG was supported by an AAPG2020-ANR-CHAIN. IM-D was supported by a National Alliance for Research on Schizophrenia and Depression (NARSAD) 2017 Young Investigator Award from the Brain & Behavior Research Foundation, the Deniker Foundation, and a 2021 Schaefer Award Scholarship from Columbia University. DJD was supported by the France 2030 programme “ANR-11-IDEX-0003,” from the OI HEALTHI of the Université Paris-Saclay, and a 2021 Schaefer Award Scholarship from Columbia University. CAD was supported by an NICHD R01 HD101402, an NIA R21 AG064774, and an NINDS R21 NS114870, and a gift from the For the Love of Travis Foundation.

BKC and CAD conceived of the study. BKC, HCH, AMG, IM-D, DJD, and CAD contributed to experimental design and intellectual interpretation. BKC wrote the manuscript. BKC, AMG, IM-D, DJD, and CAD edited the manuscript. BKC collected behavioral data. VL collected electrophysiological data. BKC, AS, MES, MP, BLW, and VP ran immunohistochemistry, processed images, and performed cell quantification. MJ performed network and statistical analysis.

We thank Dr. Jonathan Javitch, Dr. Gergely Turi, Dr. Ron Katz, and members of the laboratory for insightful comments on this project and paper. Figures of behavioral timelines were created with <u>BioRender.com</u>.

BKC, HCH, VL, AMG, IM-D, DJD, and CAD are named on provisional patent applications for the prophylactic use of (*R*,*S*)-ketamine, 5-HT_4_R agonists, and other compounds against stress-related psychiatric disorders and Alzheimer’s disease. MJ, AS, MES, MP, BLW, and VP have no conflicts of interest to disclose.

## ARTICLE INFORMATION

Address correspondence to Christine Ann Denny, Ph.D., at. MES is currently affiliated with the Icahn School of Medicine at Mount Sinai, New York, NY. VP is currently affiliated with the T.H. Chan School of Medicine at the University of Massachusetts Amherst.

## SUPPLEMENTAL INFORMATION

## SUPPLEMENTAL METHODS

### Mice

Male and female 129S6/SvEvTac mice were purchased from Taconic (Hudson, NY) at 7 weeks of age. Mice were housed 5 per cage in a 12-h (06:00-18:00) light-dark colony room at 22°C. All experiments were approved by the Institutional Animal Care and Use Committee (IACUC) at the New York Psychiatric Institute (NYSPI).

### Behavioral Assays

For all experiments, food and water were provided *ad libitum*, unless otherwise noted. Behavioral testing was performed during the light phase.

#### Contextual Fear Conditioning (CFC)

A 3-shock CFC paradigm was administered as previously described (1,2). CFC was conducted in chambers obtained from Med Associates (St. Albans, VT), with internal dimensions of approximately 20 cm wide x 16 cm deep x 20.5 cm high. The chambers had metal walls on each side, clear plastic front and back walls and ceilings, and stainless-steel bars on the floor. A house light (CM1820 bulb, 28v, 100mA) mounted directly above the chamber provided illumination. Each chamber was located inside a larger, insulated, plastic cabinet that provided protection from outside light and noise. Each cabinet contained a ventilation fan that was operated during the sessions. A paper towel dabbed with lemon solution was placed underneath the chamber floor. Mice were held outside the experimental room in their home cages prior to testing and transported to the conditioning apparatus individually in standard mouse cages. Chambers were cleaned with 70% EtOH between each set of mice. Mice were placed into the conditioning chamber and received shocks at 180 s, 240 s, and 300 s (2 s duration, 0.75 mA). Fifteen seconds after the last shock, mice were removed from the chamber. Overall, the training session lasted 317 s. During re-exposure, mice were placed in the conditioning chamber for 5 minutes and did not receive any shocks. All sessions were scored for freezing using FreezeFrame4.

#### Forced Swim Test (FST)

The FST was administered as previously described (3–5). Briefly, mice were placed into clear plastic buckets 20 cm in diameter and 23 cm deep filled 2/3 of the way with 22**°**C water. Mice were videotaped from the side for 6 min and were exposed to the swim test on 2 consecutive days. Immobility time was scored by an experimenter blind to the experimental groups.

#### Open Field (OF)

The OF assay was administered as previously described (3–5). Briefly, motor activity was quantified in 4 open field boxes 43 x 43 cm^2^ (MED Associates, Georgia, VT). An overhead camera was used to track locomotor activity. Activity chambers were computer interfaced for data sampling at 100-ms resolution. The computer defined grid lines that dividing center and surround regions, with the center square consisting of four lines 11 cm from the wall.

#### Elevated Plus Maze (EPM)

Testing was performed as previously described (3–5). Briefly, the maze is a plus-cross-shaped apparatus consisting of four arms, two open and two enclosed by walls, linked by a central platform at a height of 50 cm from the floor. Mice were individually placed in the center of the maze facing an open arm and were allowed to explore the maze for 5 min. The time spent in and the number of entries into the open arms was used as an anxiety index. Videos were scored using ANY-maze behavior tracking software (Stoelting, Wood Dale, IL).

#### Marble Burying (MB)

The MB assay was conducted in a clean cage (10.5 in x 5.5 in) containing soft pliable Beta Chip bedding (Northeastern Products Corp, Warrensburg, NY). The cage contained 16 marbles set up in 4 rows of 4 across. Mice were given 30 minutes to explore and bury. At the end of the assay, the percentage of marbles buried was calculated.

#### Novelty Suppressed Feeding (NSF)

Testing was performed as previously described (4,5). Briefly, the NSF testing apparatus consisted of a plastic box (50 x 50 x 20 cm). The floor of which was covered with approximately 2 cm of wooden bedding and the arena was brightly lit (approximately 1000 lux). Mice were food restricted for 12 h prior to testing. At the time of testing, a single pellet of food (regular chow) was placed on a white paper platform positioned in the center of the box. Each animal was placed in a corner of the box, and a stopwatch was immediately started. The latency of the mice to begin eating in the arena was recorded. Immediately after the latency was recorded, the food pellet was removed from the arena. The mice were then placed back into their home cage. The latency to eat and the amount of food consumed in 5 min were measured (home cage consumption), followed by an assessment of post-restriction weight. A Kaplan-Meier survival analysis was used due to the lack of normal distribution of data. The Mantel-Cox log-rank test was used to evaluate differences between the experimental groups.

### Electrophysiology

Electrophysiology was conducted as previously described (5). One week after a saline or a (*R,S*)-ketamine (10 mg/kg) + prucalopride (3 mg/kg) injection, mice were anesthetized by isoflurane inhalation, decapitated, and brains rapidly removed. CA3 slices (350 μm) were cut on a vibratome (Leica VT1000S) in ice cold partial sucrose artificial cerebrospinal fluid (ACSF) solution (in mM): 80 NaCl, 3.5 KCl, 4.5 MgSO_4_, 0.5 CaCl_2_, 1.25 H_2_PO4, 25 NaHCO_3_, 10 C_6_H_12_O_6_, and 90 C_12_H_22_O_11_ equilibrated with 95% O_2_ / 5% CO_2_ and stored in the same solution at 37°C for 30 minutes, then at room temperature until use. Recordings were made at 30-32°C (TC324-B; Warner Instrument Corp) in ACSF (in mM: 124 NaCl, 8.5 KCl, 1 NaH_2_PO_4_, 25 NaHCO_3_, 20 glucose, 1 MgCl_2_, 2 CaCl_2_). Whole-cell voltage clamp recordings (-70 mV) were obtained using a patch pipette (4-6 M MΩ) containing (in mM): 135 K Gluconate, 5 KCl, 0.1 EGTA-Na, 10 HEPES, 2 NaCl, 5 ATP, 0.4 GTP, 10 phosphocreatine (pH 7.2; 280–290 mOsm). Bicuculline (5 μM) was also included in the bath solution to inhibit GABA_A_ receptors. Three-dihydroxy-6-nitro-7-sulfamoyl-benzo[f]quinoxaline (NBQX) (20 mM) was added later in recordings to inhibit AMPAR synaptic currents and unmask NMDAR-mediated signals. Patch pipettes were made from borosilicate glass (A-M Systems, Sequium, WA) using a micropipette puller (Model P-1000; Sutter Instruments, Novato, CA).

Recordings were made without correction for junction potentials. Pyramidal cells were visualized and targeted via infrared-differential interference contrast (IR-DIC; 40x objective) optics on an Axioskop-2 FS (Zeiss, Oberkochen, Germany).

### Immunohistochemistry

Immunohistochemistry was performed as previously described (2,5). Mice were deeply anesthetized, and brains were fixed and extracted using transcardial perfusion. For c-fos immunohistochemistry, floating sections were used. Sections were first rinsed 3 times in 1 x phosphate buffered saline (PBS) and then blocked in 1 x PBS with 0.5% Triton X-100 (PBST) and 10% normal donkey serum (NDS) for 2 hours at room temperature (RT). Incubation with primary antibodies was performed at 4°C overnight (rat anti-c-fos, 226 017, 1:5000, SySy, Göettingen, Germany; rabbit anti-parvalbumin, PV27, 1:3000, Swant, Burgdorf, Switzerland) in 1 x PBST. Sections were then washed 3 times in 1 x PBS and incubated with secondary antibody (Alexa 647 anti-rat, Ab150155, 1:500, Abcam, Cambridge, UK; Alexa 488 anti-rabbit, A-21206, 1:500, Thermo Fisher Scientific, Waltham, MA) for 2 hours at RT. Sections were then washed three times in 1 x PBS, mounted on slides, and coverslipped with Fluoromount G (Electron Microscopy Sciences, Hatfield, PA).

### Confocal Microscopy

Fluorescent confocal micrographs were captured with a Leica TCS SPE-II confocal microscope with LAS X software as previously described (4,5). Bilateral hippocampal sections were imaged throughout the rostro-caudal axis of the HPC using a 20X objective. Identification of hippocampal regions involved acquiring 6 dorsal and ventral sections per mouse brain slice at 20X. All individual panels were acquired at a thickness of 3 μm. Z-stack analysis was performed using the LAS X image browser to determine expression of c-fos and PV. Expression levels of c-fos and PV were compared across all sections using identical exposure conditions.

### Cell Quantification

An investigator blind to treatment groups used Fiji software to count c-fos^+^, PV^+^, and c-fos^+^/PV^+^ immunoreactive cells in the ACA, ILA, and PL of the mPFC or in the DG, CA3, and CA1 throughout the entire rostrocaudal axis of the HPC. Cells were counted bilaterally. Number of c-fos^+^, PV^+^, and c-fos^+^/PV^+^ cells is presented throughout the text.

### Correlation and Network Analyses

Regions were mapped across all experimental groups and included a minimum n of 5 mice per group. Pearson correlations between regions were calculated using the Hmisc package in R, with pairwise removal of missing cases. Significance of pairwise regional correlation differences between experimental groups was calculated using permutation analysis. Group labels were randomly shuffled, and correlations were recomputed 1000 times to generate a null distribution of the pairwise regional correlation differences between groups. Correlation differences were compared to these null distributions to determine the p-value. For the cluster network maps, relevant functional connections were retained by thresholding connections at p < 0.05. To ensure that both relevant positive and negative functional connections were used for community detection, the absolute Pearson values were used as edge weights. Using the igraph and tidygraph packages, the cluster fast greedy algorithm was used for community detection and visualized as a color-coded force-directed network (Fruchterman and Reingold layout) and as a dendrogram. Communities and nodes were color-coded, and scales were changed for edge connections (correlation strength).

To compare global network properties of c-Fos^+^ and PV^+^ expression across experimental groups, networks were again constructed based on Pearson correlations and edges were thresholded at a p < 0.05. For each region, the clustering coefficient and measures of centrality, such as degree, betweenness centrality, and efficiency were calculated using the tidygraph and igraph packages. These measures were averaged across all regions to calculate global network statistics. Networks were visualized using ggraph and summary statistics were plotted in Prism 10 (Graphpad Software, La Jolla, CA).

### Statistical Analysis

Data were analyzed using Prism 9.0 (Graphpad Software, La Jolla, CA). Jmp 16 was used for bootstrap prediction analysis. Alpha was set to 0.05 for all analyses. Generally, the effect of Drug or Group was analyzed using an analysis of variance (ANOVA), using repeated measures where appropriate. Post-hoc Dunnett, Sidak, or Tukey tests were used where appropriate. All statistical tests and *p* values are listed in **Tables S01 and S02**.

## SUPPLEMENTAL FIGURES AND FIGURE LEGENDS

**Figure S1.**
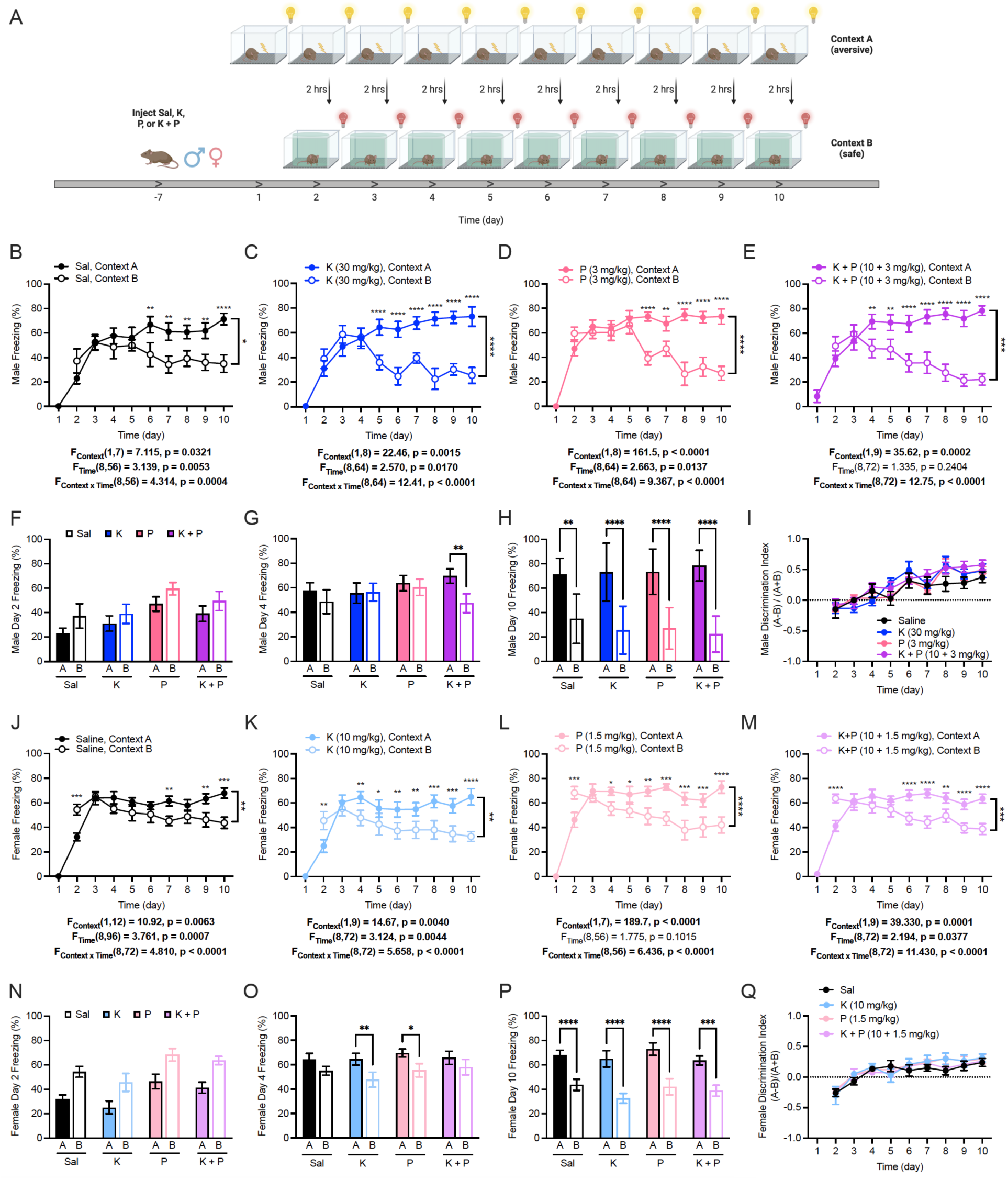
Combined prophylactic (*R,S*)-ketamine + prucalopride administration allows for faster contextual fear discrimination in male mice. **(A)** Experimental design. Female and male mice were administered Sal, K (30 mg/kg in male mice, 10 mg/kg in female mice), P (3 mg/kg in male mice, 1.5 mg/kg in female mice), or K+P (10 + 3 mg/kg in male mice, 10 + 1.5 mg/kg in female mice) one week prior to the start of CFD. **(B)** Sal-administered male mice started discriminating on day 6. **(C)** K-administered male mice started discriminating on day 5. **(D)** P-administered male mice started discriminating on day 6. **(E)** K+P-administered male mice started discriminating on day 4. Average freezing on days **(F)** 2, **(G)** 4, and **(H)** 10 for male mice. **(I)** All groups of male mice showed comparable discrimination ratios during CFD. **(J)** Sal-administered female mice started discriminating on day 7. **(K)** K-administered female mice started discriminating on day 4. **(L)** P-administered female mice started discriminating on day 4. **(M)** K+P-administered female mice started discriminating on day 6. Average freezing on days **(N)** 2, **(O)** 6, and **(P)** 10 for female mice. **(Q)** All groups of female mice showed comparable discrimination ratios during CFD. (n = 6-10 female mice per group). Error bars represent ± SEM. * p < 0.05. ** p < 0.01. *** p < 0. 001, **** p < 0.0001. Sal, saline; K, (*R,S*)-ketamine; P, prucalopride; K + P, (*R,S*)-ketamine + prucalopride; hrs, hours; mg, milligram; kg, kilogram.

**Figure S2.**
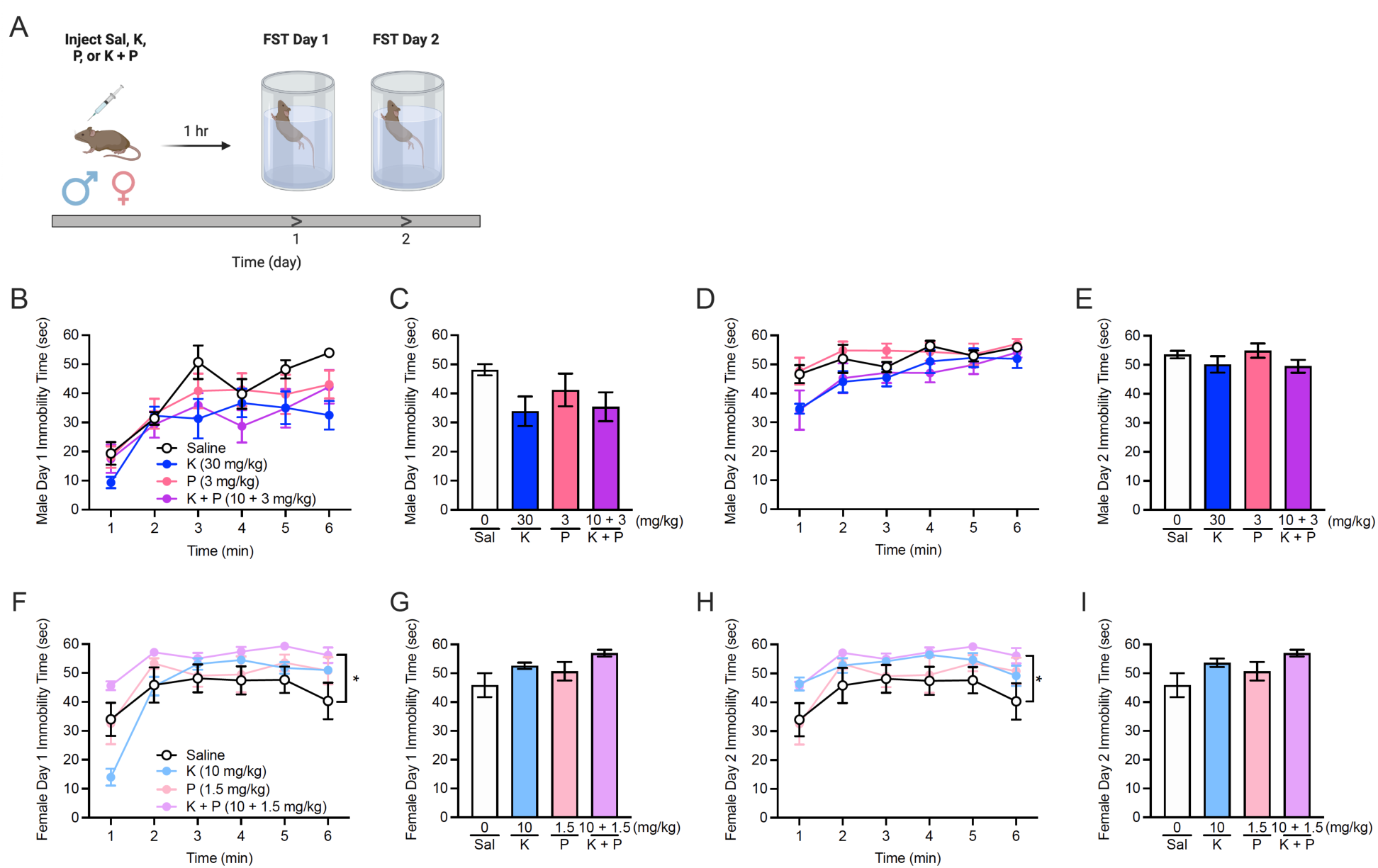
Combined (*R,S*)-ketamine + prucalopride does not attenuate immobility time in the FST in non-stressed 129S6/SvEv mice. **(A)** Experimental protocol. Sal, K, P, or K+P was administered to male or female mice one hour prior to the FST. In male mice, immobility time was comparable during **(B-C)** day 1 and **(D-E)** day 2 of the FST. **(F-G)** On days 1 and 2 of the FST, K+P-administered female mice exhibited higher overall immobility time when compared to saline controls, but this effect was not significant when comparing average immobility. (n = 4-7 mice per group). Error bars represent ± SEM. * p < 0.05. Sal, saline; K, (*R,S*)-ketamine; P, prucalopride; FST, forced swim test; hr, hour; sec, seconds; min, minutes; mg, milligrams; kg, kilograms.

**Figure S3.**
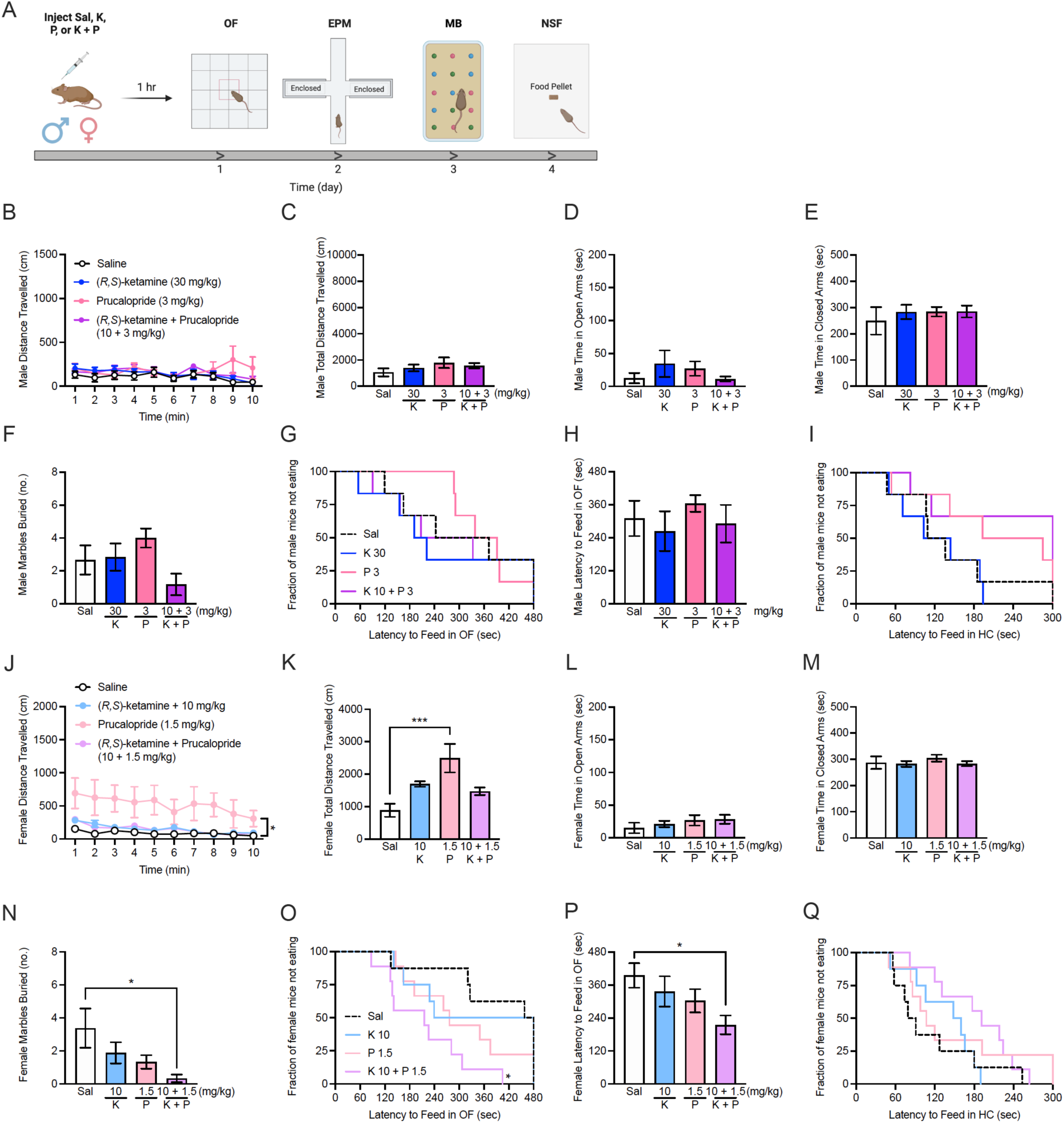
Combined (*R,S*)-ketamine + prucalopride reduces perseverative behavior in non-stressed female mice. **(A)** Experimental protocol. Male or female 129S6/SvEv mice were given a single injection of Sal, K, P, or K+P one hour prior to the OF test. On subsequent days, mice were administered the EPM, MB, and NSF assays. **(B-I)** Behavior in the OF, EPM, MB, and NSF assays was comparable across all groups of male mice. **(J-K)** Distance traveled in the OF was significantly higher in prucalopride-administered female mice. **(L-M)** Time spent in the open and closed arms of the EPM was comparable in all groups of female mice. **(N)** Female mice given P or K+P buried a lower number of marbles when compared to Sal-administered mice. **(O-Q)** Behavior in the NSF in female mice was comparable across all groups. Error bars represent ± SEM. * p < 0.05. Sal, saline; K, (*R,S*)-ketamine; P, prucalopride; OF, open field; h, hour; EPM, elevated plus maze; MB, marble burying; NSF, novelty-suppressed feeding; cm, centimeters; min, minutes; mg, milligrams; kg, kilograms; sec, seconds; no., number.

**Figure S4.**
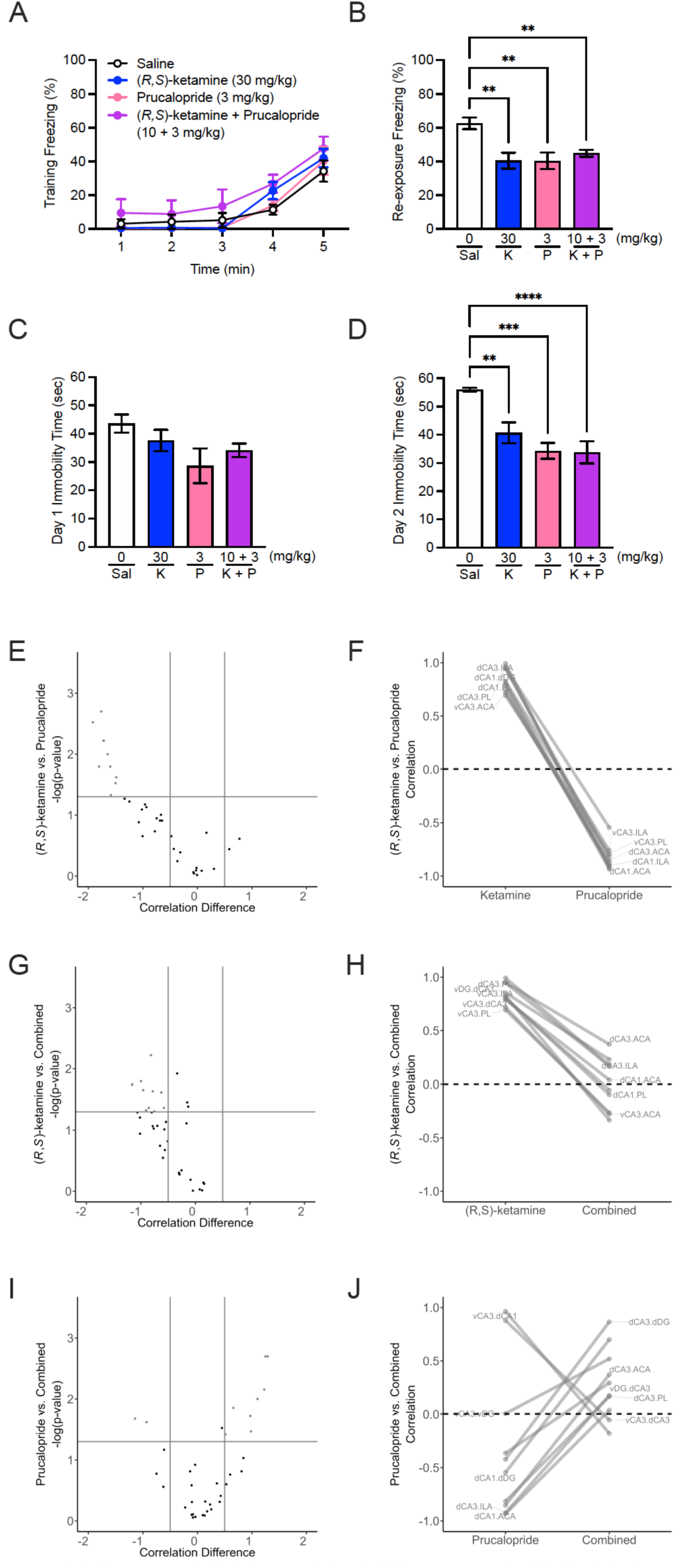
Combined (*R,S*)-ketamine + prucalopride alters correlated excitatory signaling in mPFC and HPC. **(A-B)** K, P, and K+P reduced freezing during CFC re-exposure but did not alter freezing during CFC training. **(C-D)** K, P, and K+P reduced immobility time during day 2, but not day 1 of the FST. Volcano and parallel plots indicating regional c-fos correlation differences greater than 0.5 between **(E-F)** K and P, **(G-H)** K and K+P, and **(I-J)** P and K+P. (n = 7-8 mice per group). Error bars represent ± SEM. ** p < 0.01, *** p < 0. 001, **** p < 0.0001. Sal, saline; K, (*R,S*)-ketamine; P, prucalopride; K + P, (*R,S*)-ketamine + prucalopride; ACA, anterior cingulate area; ILA, infralimbic area; PL, prelimbic area; dDG, dorsal dentate gyrus; dCA3, dorsal field CA3; dCA1, dorsal field CA1; vDG, ventral dentate gyrus; vCA3, ventral field CA3; vCA1, ventral field CA1.

**Figure S5.**
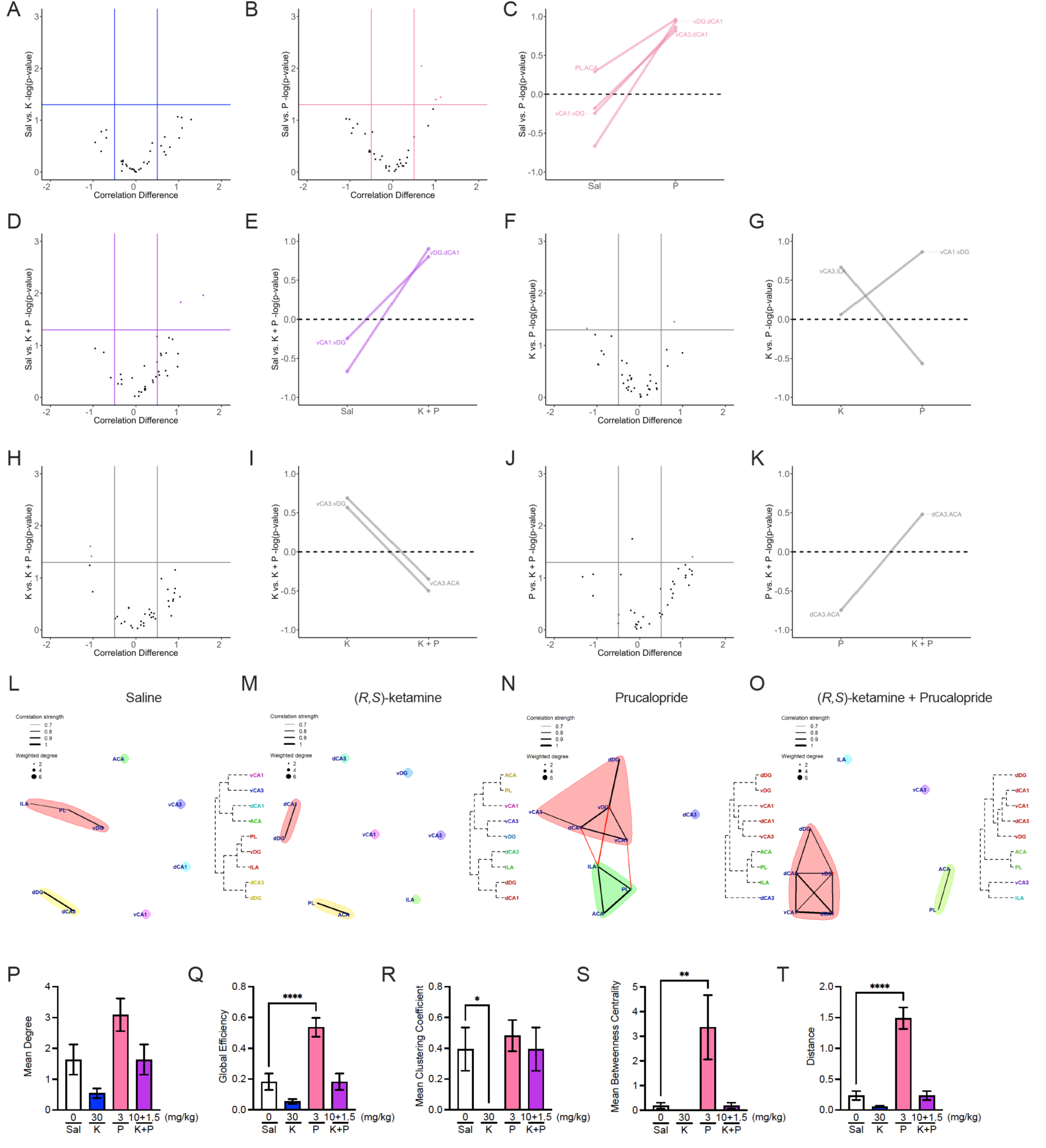
Combined (*R,S*)-ketamine + prucalopride alters correlated PV expression and inhibitory network activity. Volcano and parallel plots indicating regional PV correlation differences greater than 0.5 between. **(A)** Sal and K, **(B-C)** Sal and P, **(D-E)** Sal and K+P, **(F-G)** K and P, **(H-I)** K and K+P, and **(J-K)** P and K+P. Notably, mice given K+P exhibited an increase in correlated vCA1-vDG and vDG-dCA1 activity when compared with Sal controls. Cluster maps of correlated PV expression in **(L)** Sal-, **(M)** K-, **(O)** P, and **(P)** K+P-administered mice reveal sparsely connected activity in Sal-and K-administered mice, while cluster maps for prucalopride and combined drug-administered cluster maps are more cohesive. In particular, vCA3 is an isolated region in the combined drug cluster map, in contrast to all other dorsal and ventral hippocampal regions. **(P)** Mean degree is comparable across all drug groups. **(Q)** P administration increases mean global efficiency when compared with saline administration. **(R)** K administration reduces the mean clustering coefficient compared to saline. **(S-T)** Mean betweenness centrality and network distance are increased in prucalopride-administered mice when compared with saline controls. (n = 7-8 mice per group). Error bars represent ± SEM. ACA, anterior cingulate area; ILA, infralimbic area; PL, prelimbic area; dDG, dorsal dentate gyrus; dCA3, dorsal field CA3; dCA1, dorsal field CA1; vDG, ventral dentate gyrus; vCA3, ventral field CA3; vCA1, ventral field CA1. Sal, saline; K, (*R,S*)-ketamine; P, prucalopride; K + P, (*R,S*)-ketamine + prucalopride; mg, milligram; kg, kilogram.

**Figure S6.**
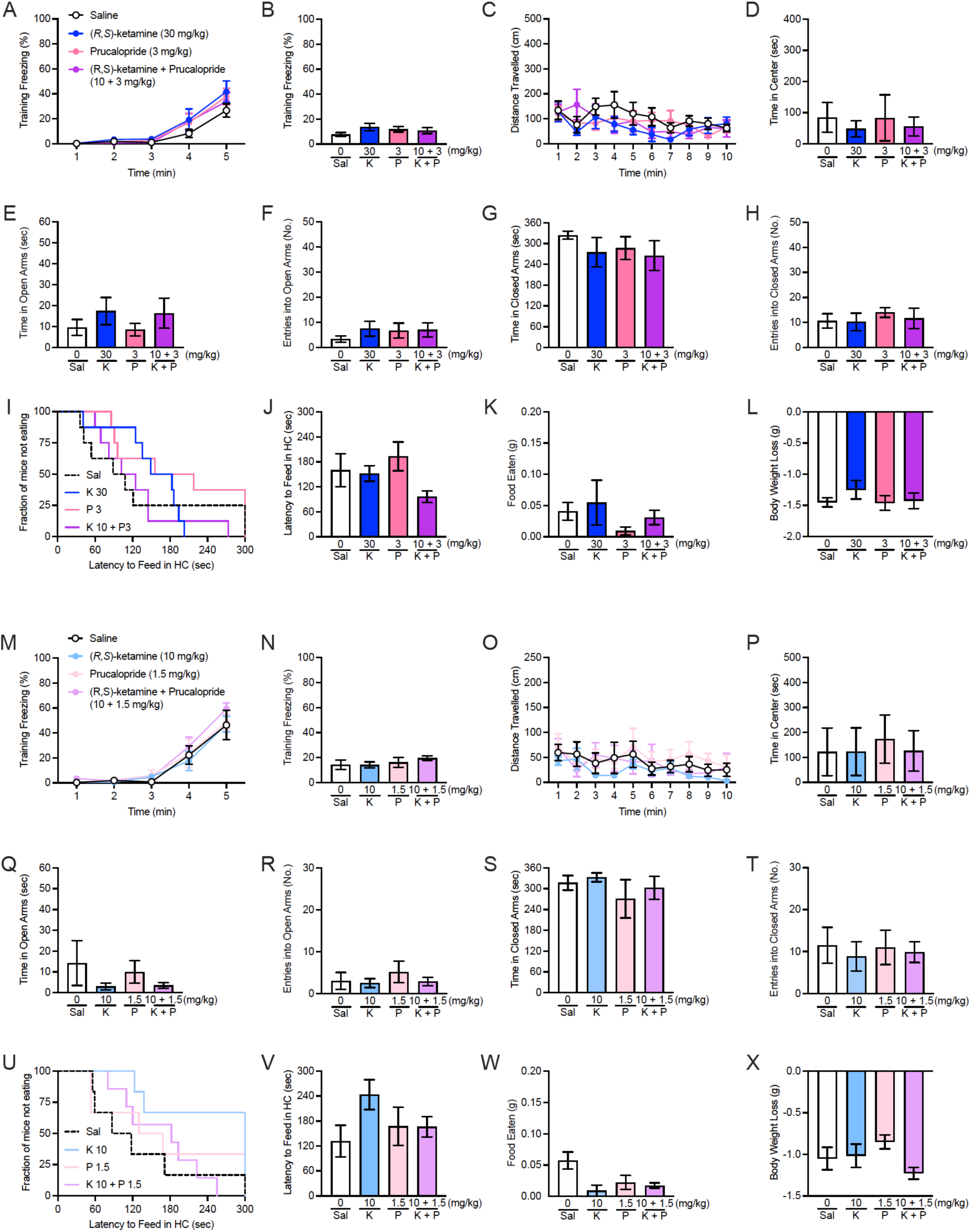
Combined (*R,S*)-ketamine + prucalopride attenuates learned fear in male mice and reduces behavioral despair and hyponeophagia in both sexes when administered after stress. **(A)** Freezing was comparable across all groups of male mice during CFC training. Behavior was comparable across all groups of male mice in **(C-D)** the OF and **(E-H)** the EPM. **(I-J)** Latency to feed in the home cage, **(K)** food eaten, and **(L)** body weight loss in the NSF were comparable across all groups of male mice. **(M-N)** Freezing during CFC training was comparable across all drug groups in female mice. In female mice, behavior in **(O-P)** the OF and **(Q-T)** the EPM was not altered by prophylactic drug administration. **(U-V)** Latency to feed in the home cage, **(W)** food eaten, and **(X)** body weight loss in the NSF were comparable across all groups of female mice. (n = 5-7 male or female mice per group). Error bars represent ± SEM. Sal, saline; K, (*R,S*)-ketamine; P, prucalopride; K + P, (*R,S*)-ketamine + prucalopride; mg, milligram; kg, kilogram; cm, centimeter; min, minute; sec, second; no., number; HC, home cage; g, gram.

**Table S01.**
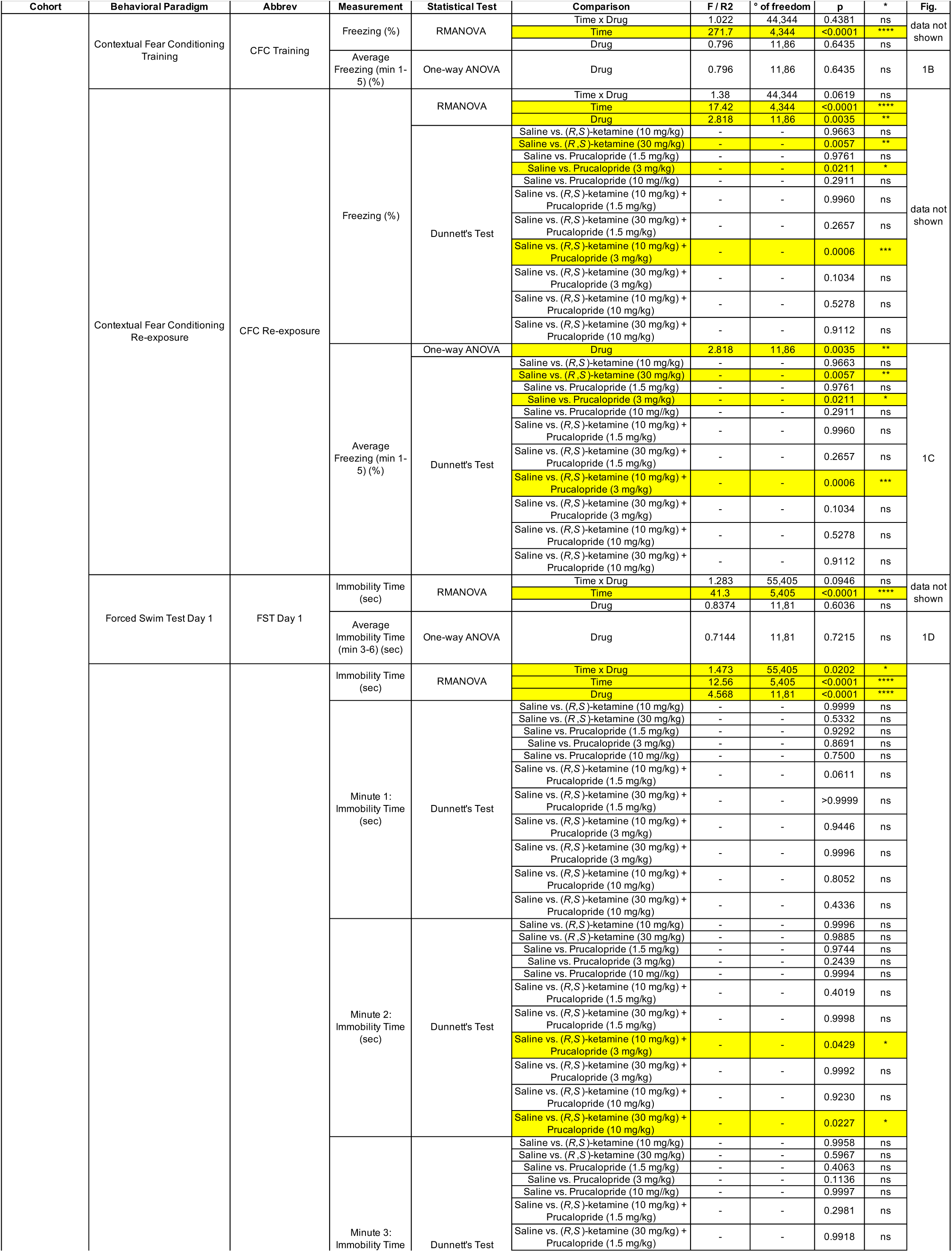
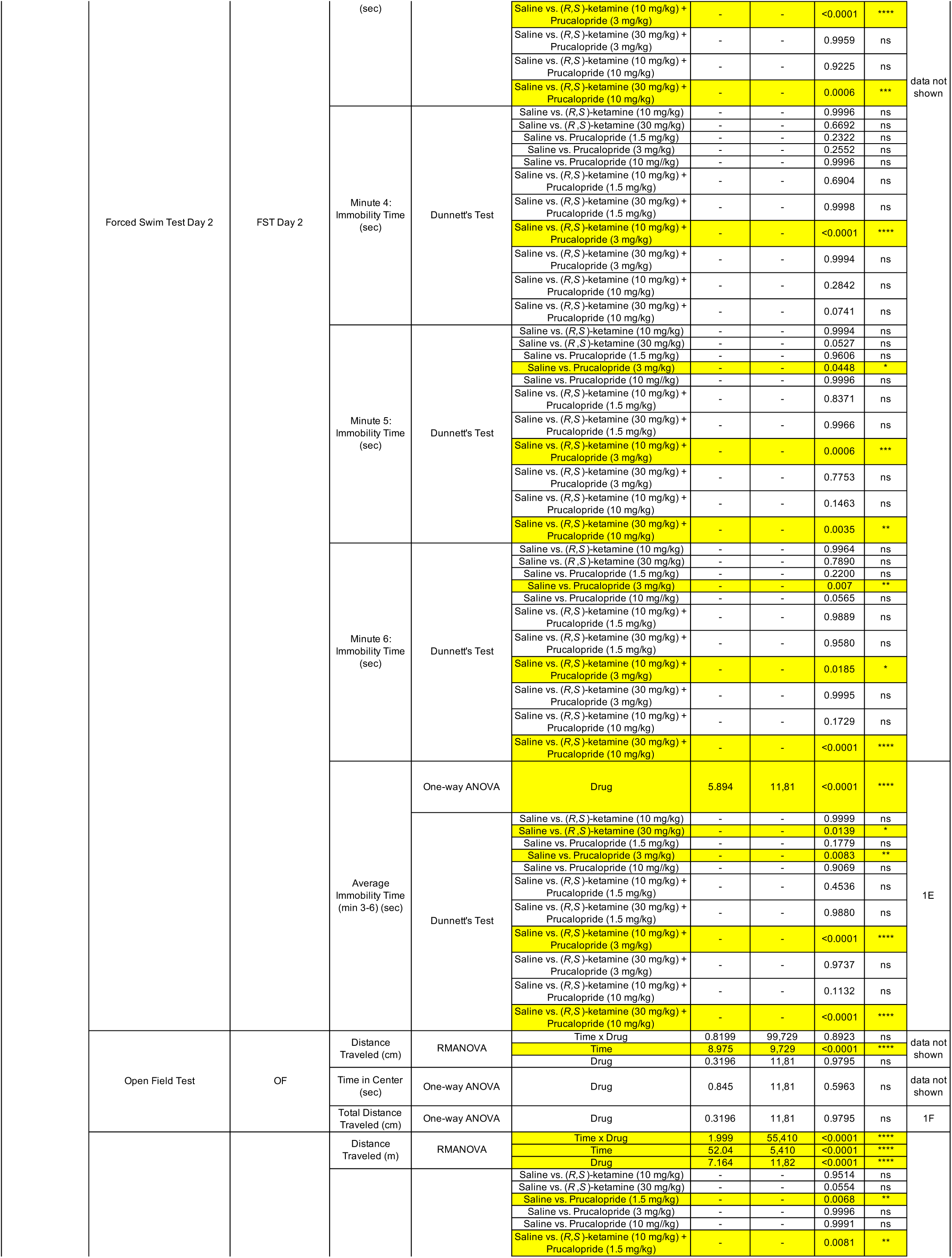
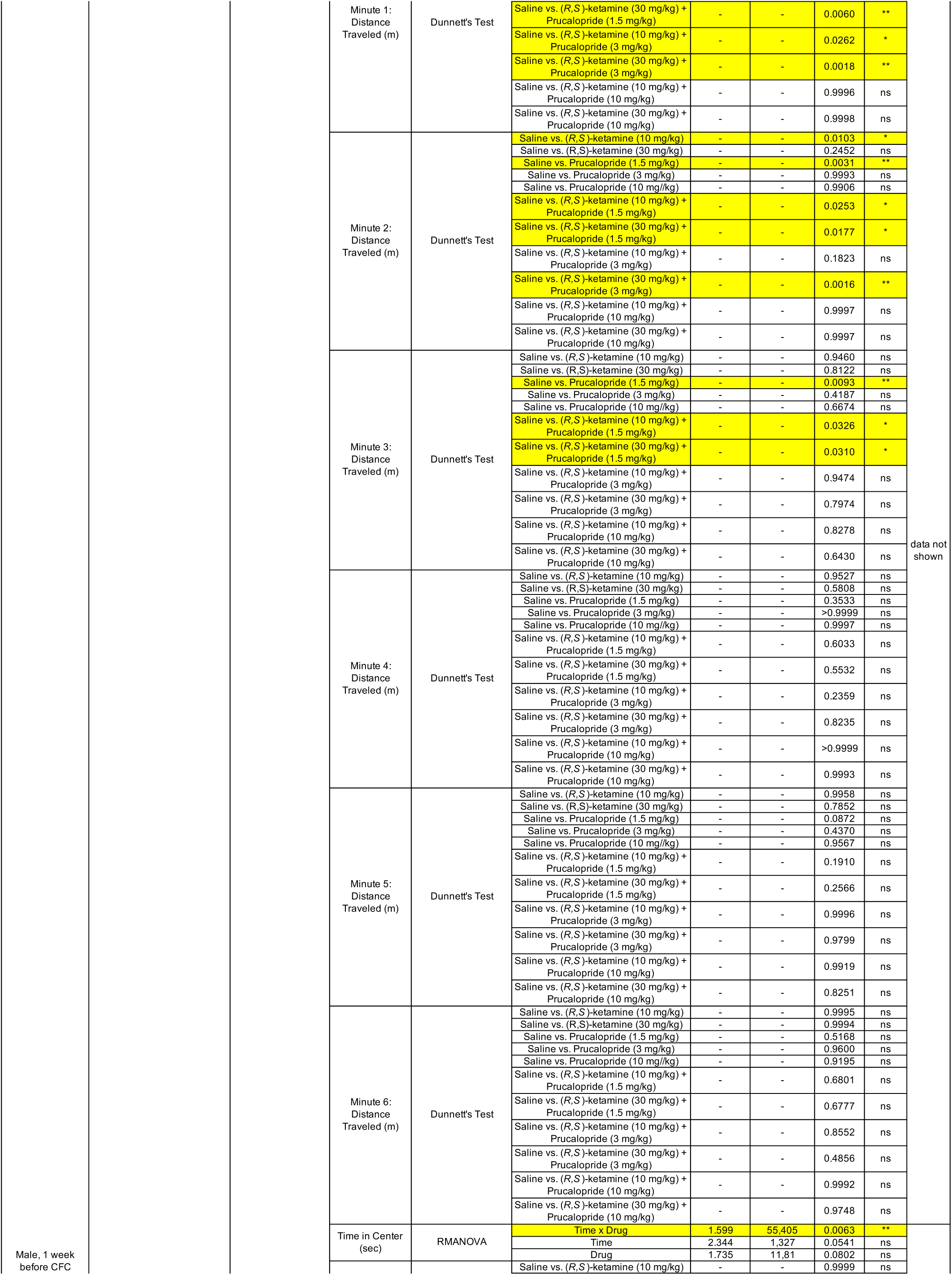
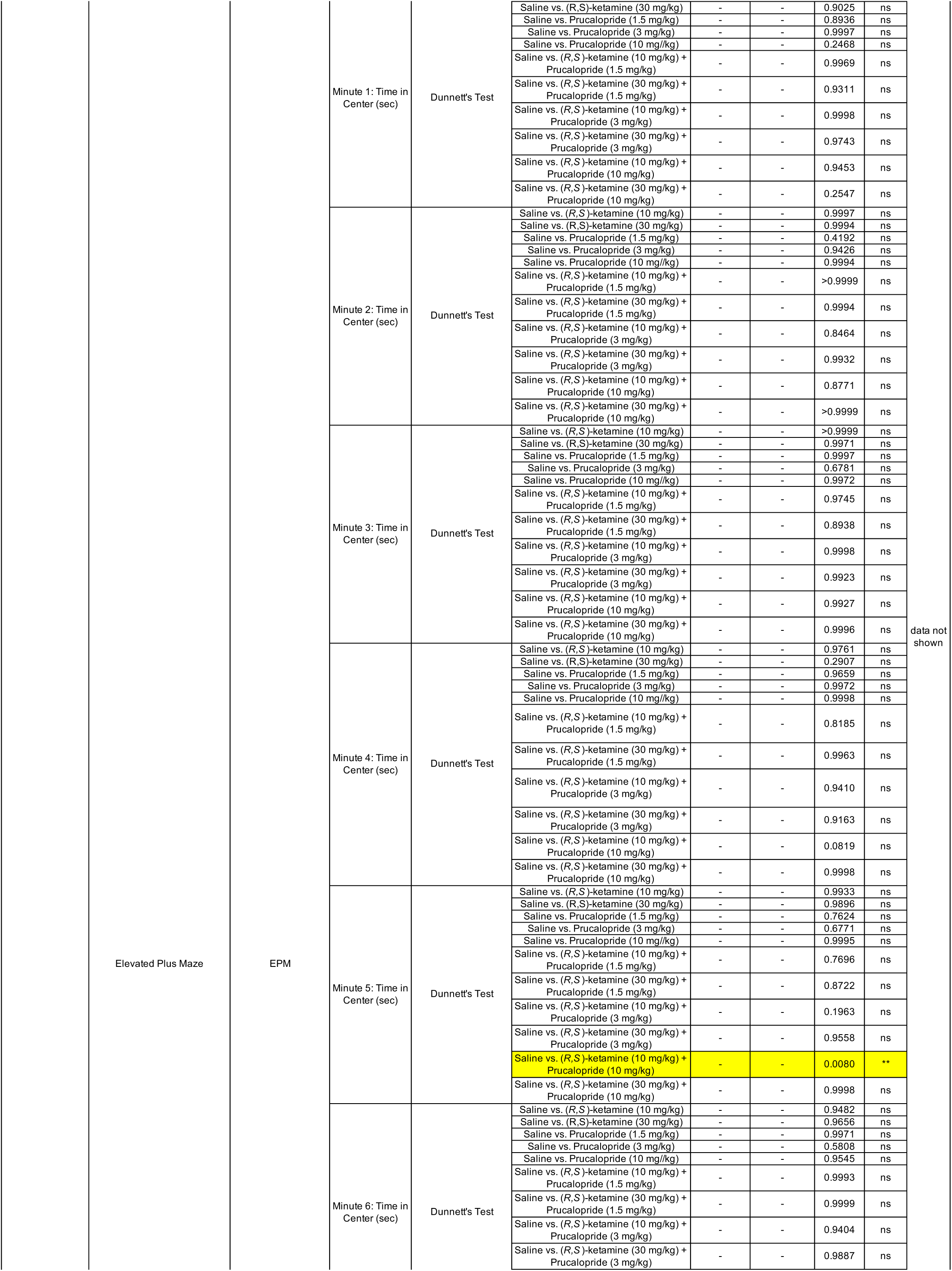
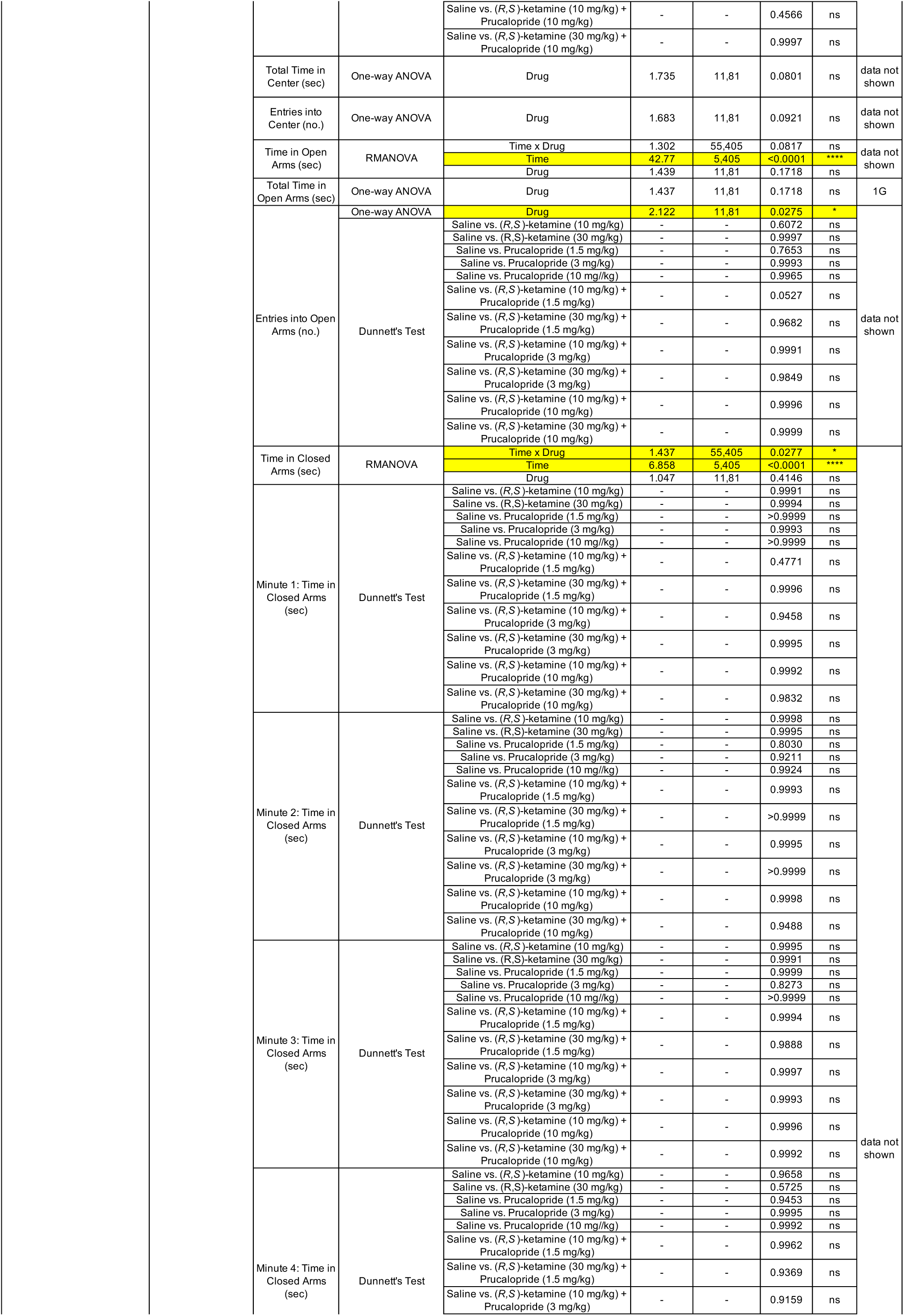
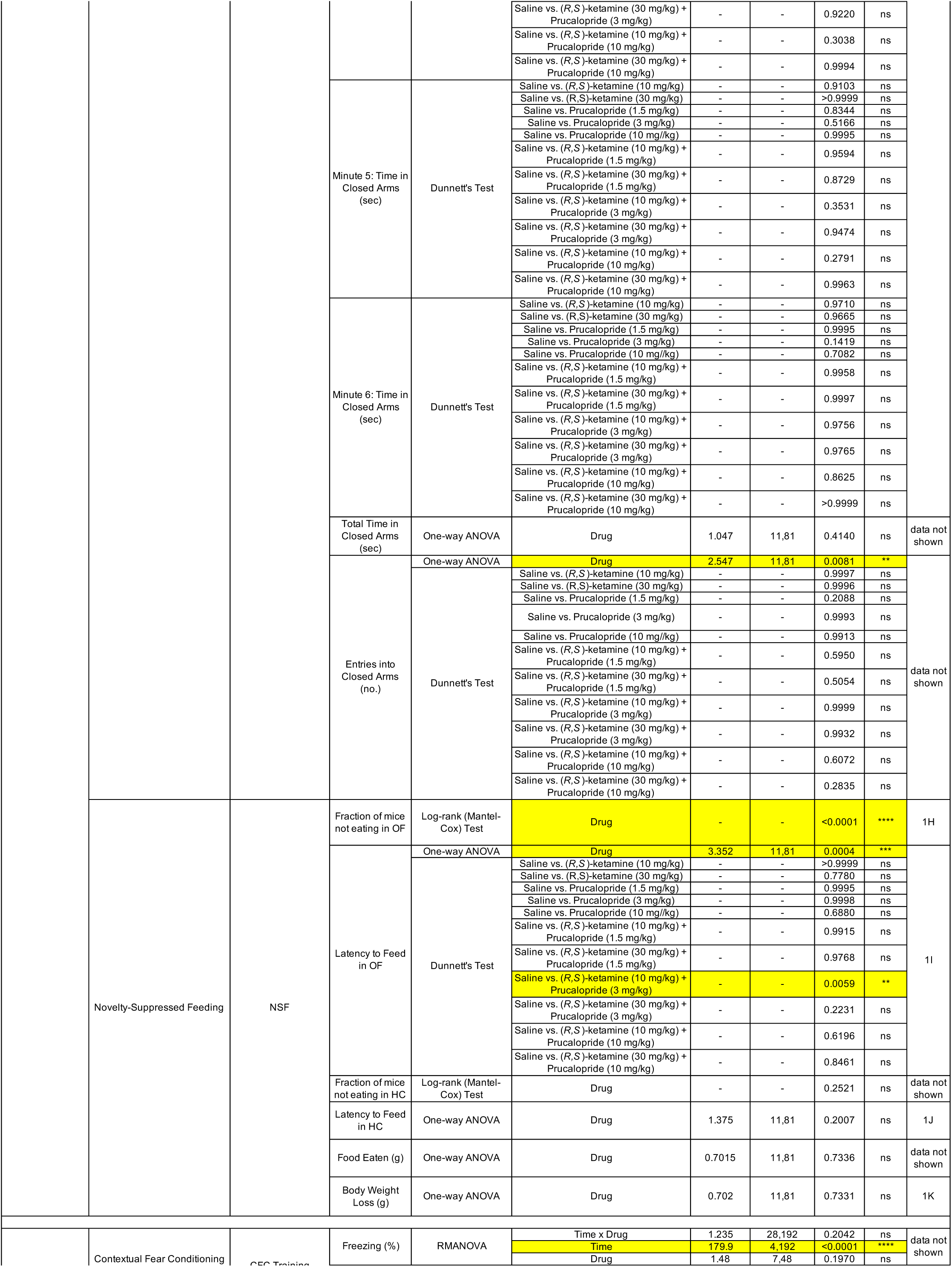
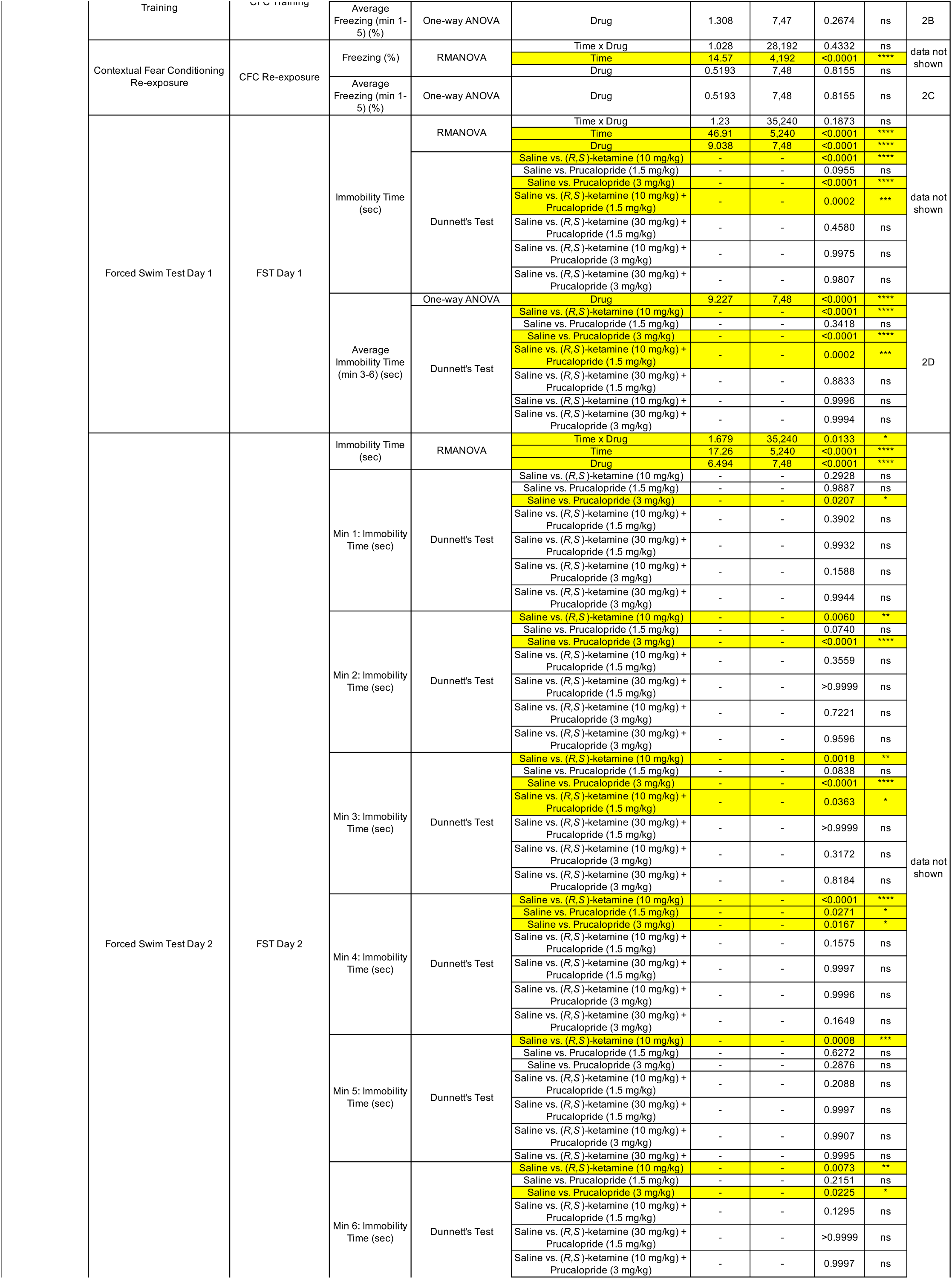
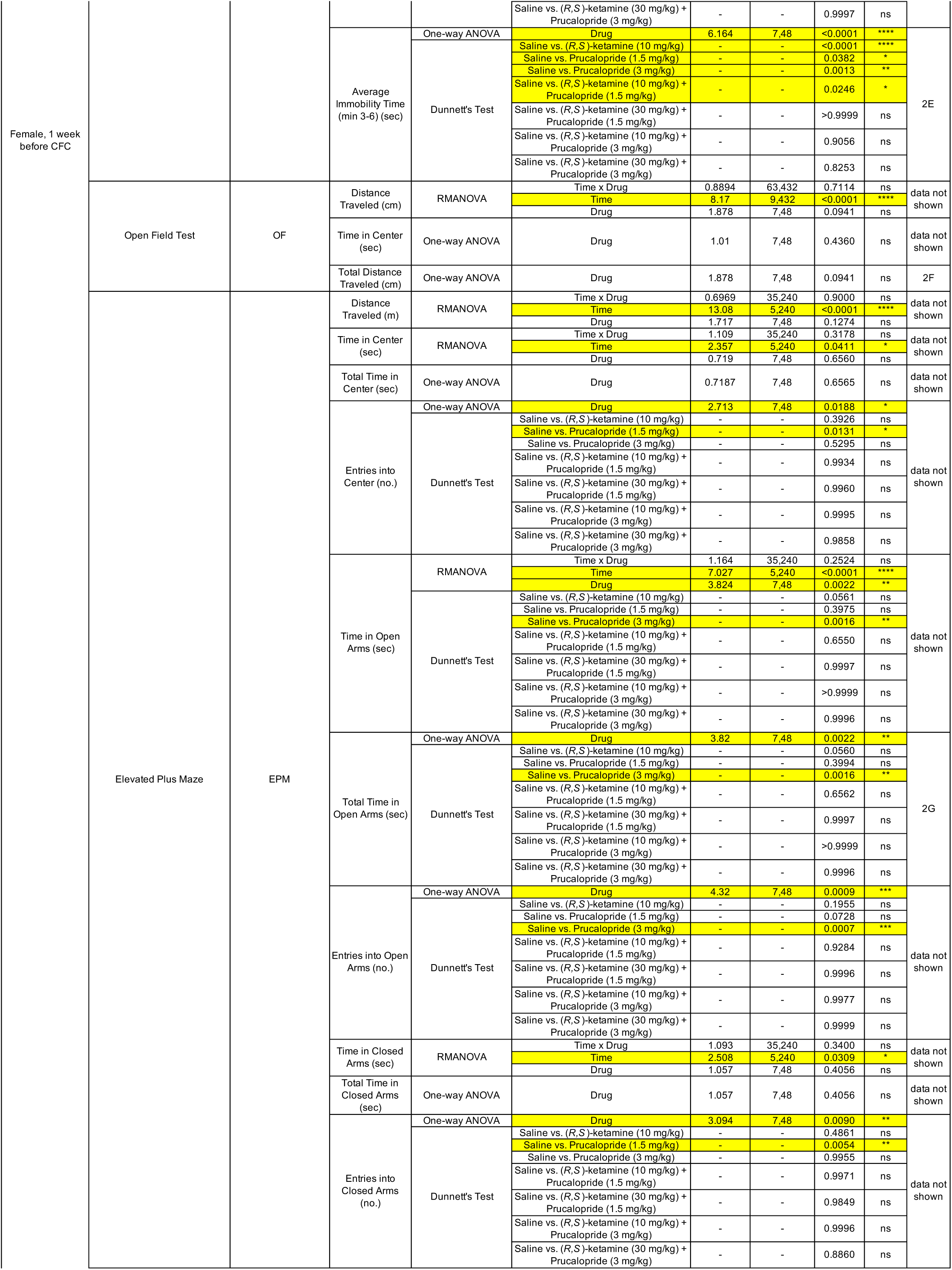
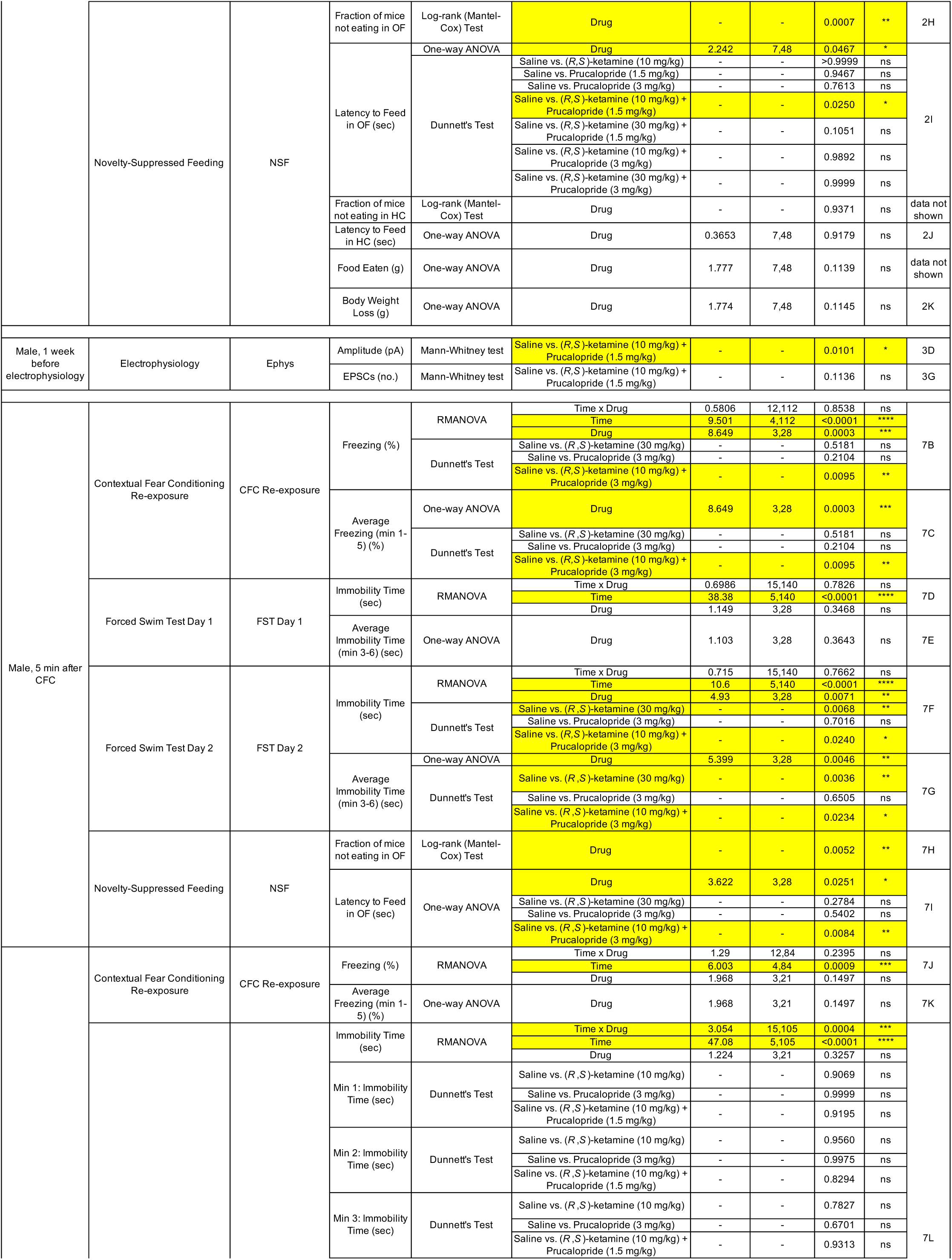
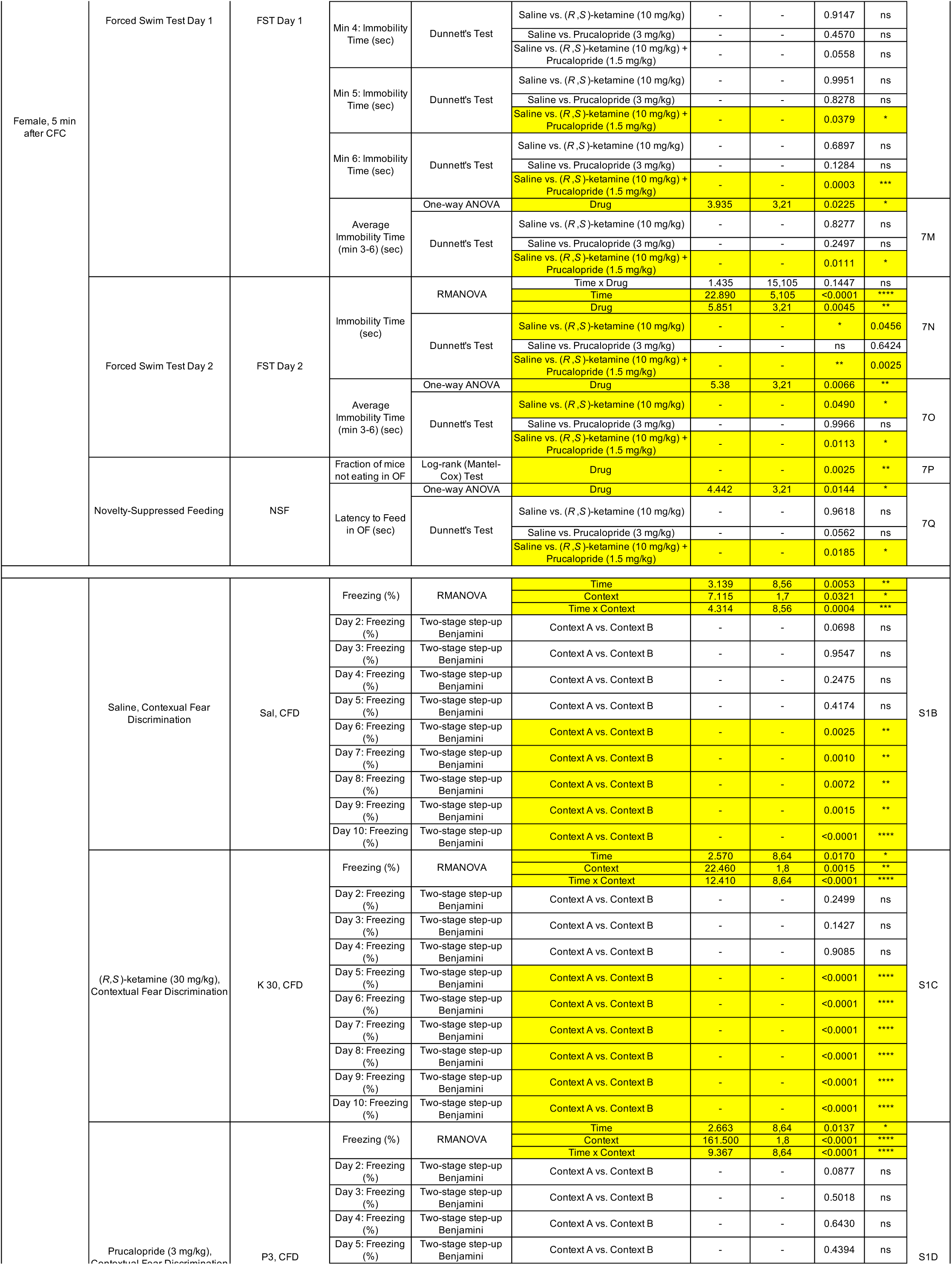
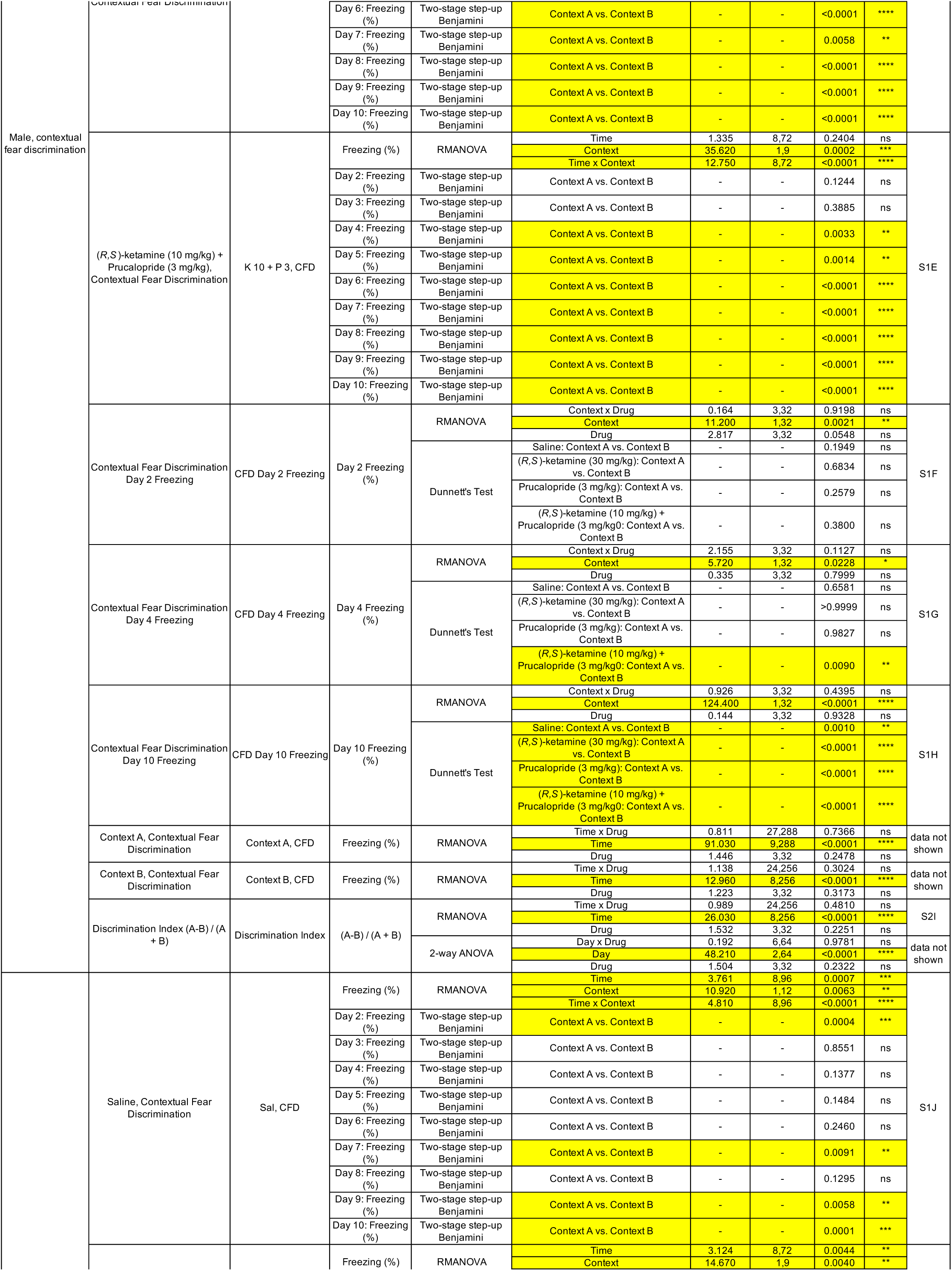
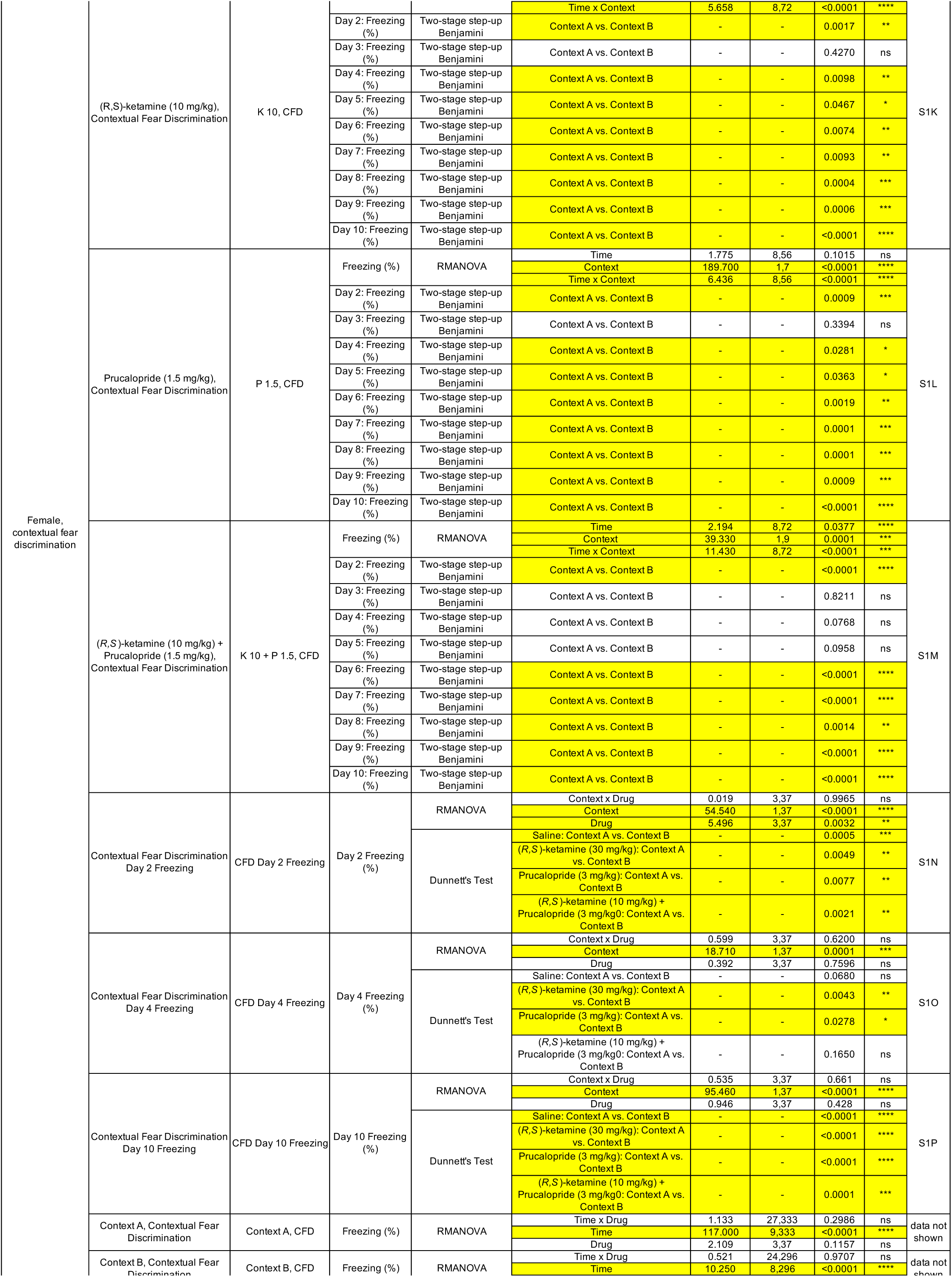
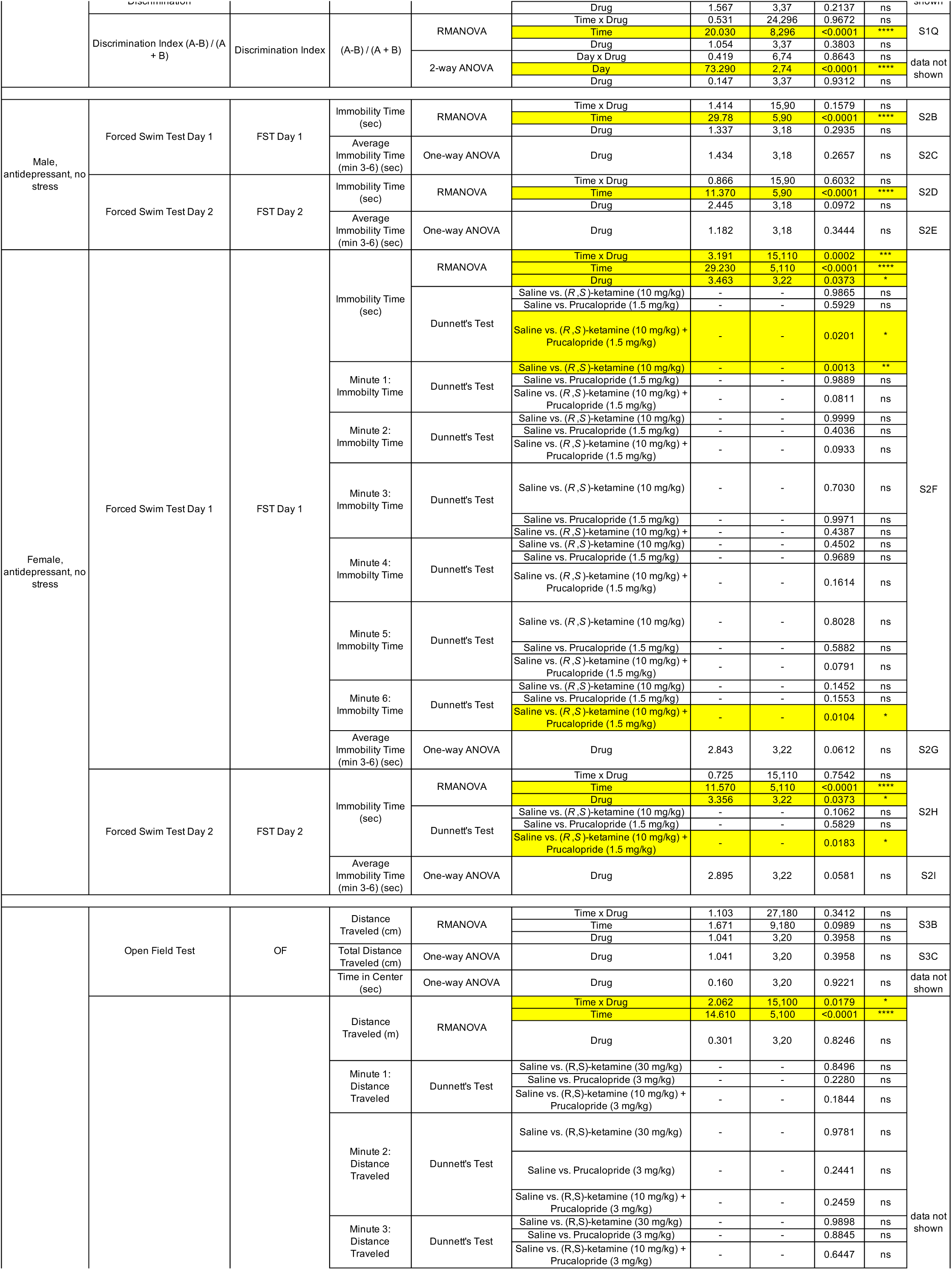
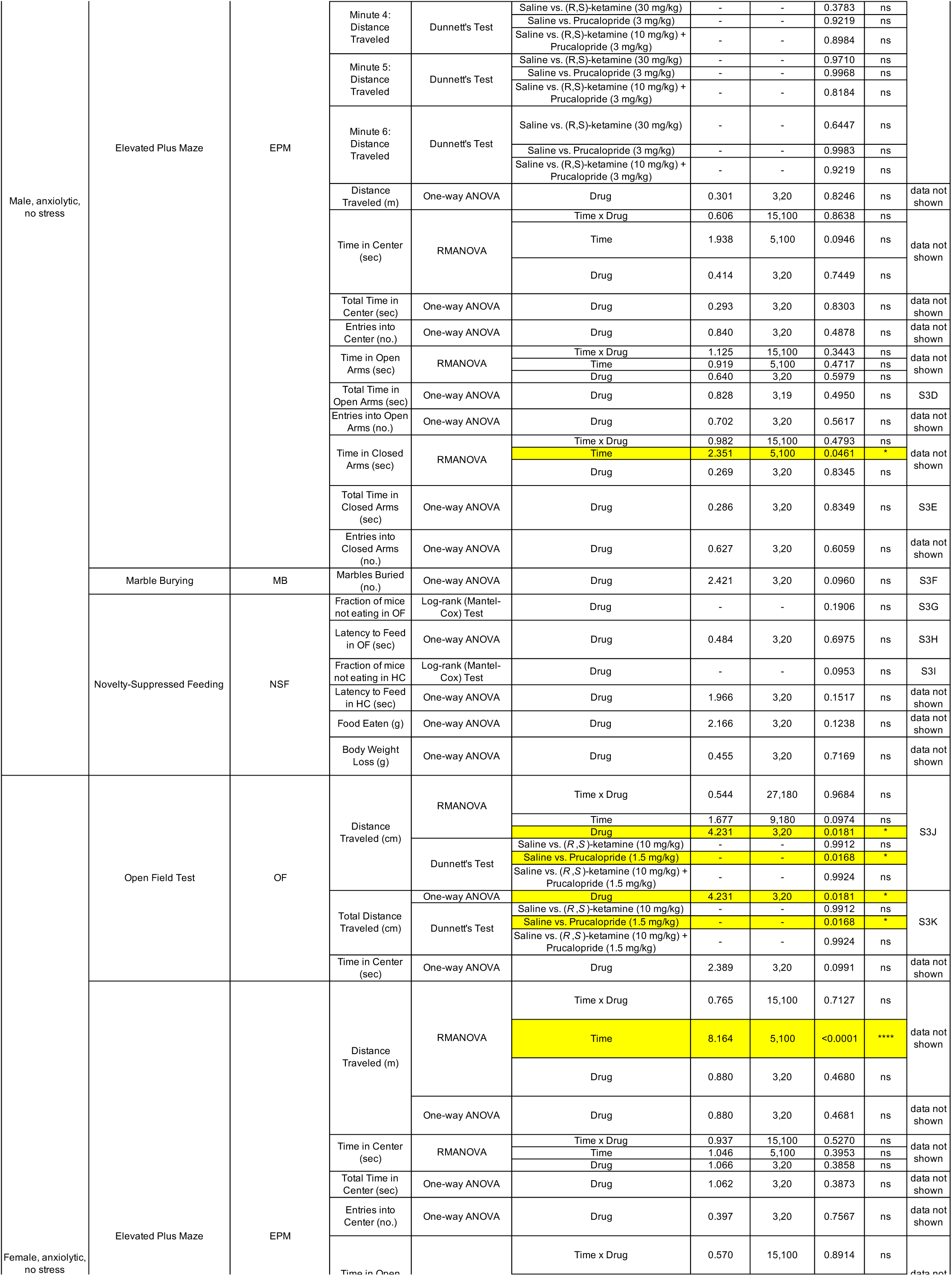
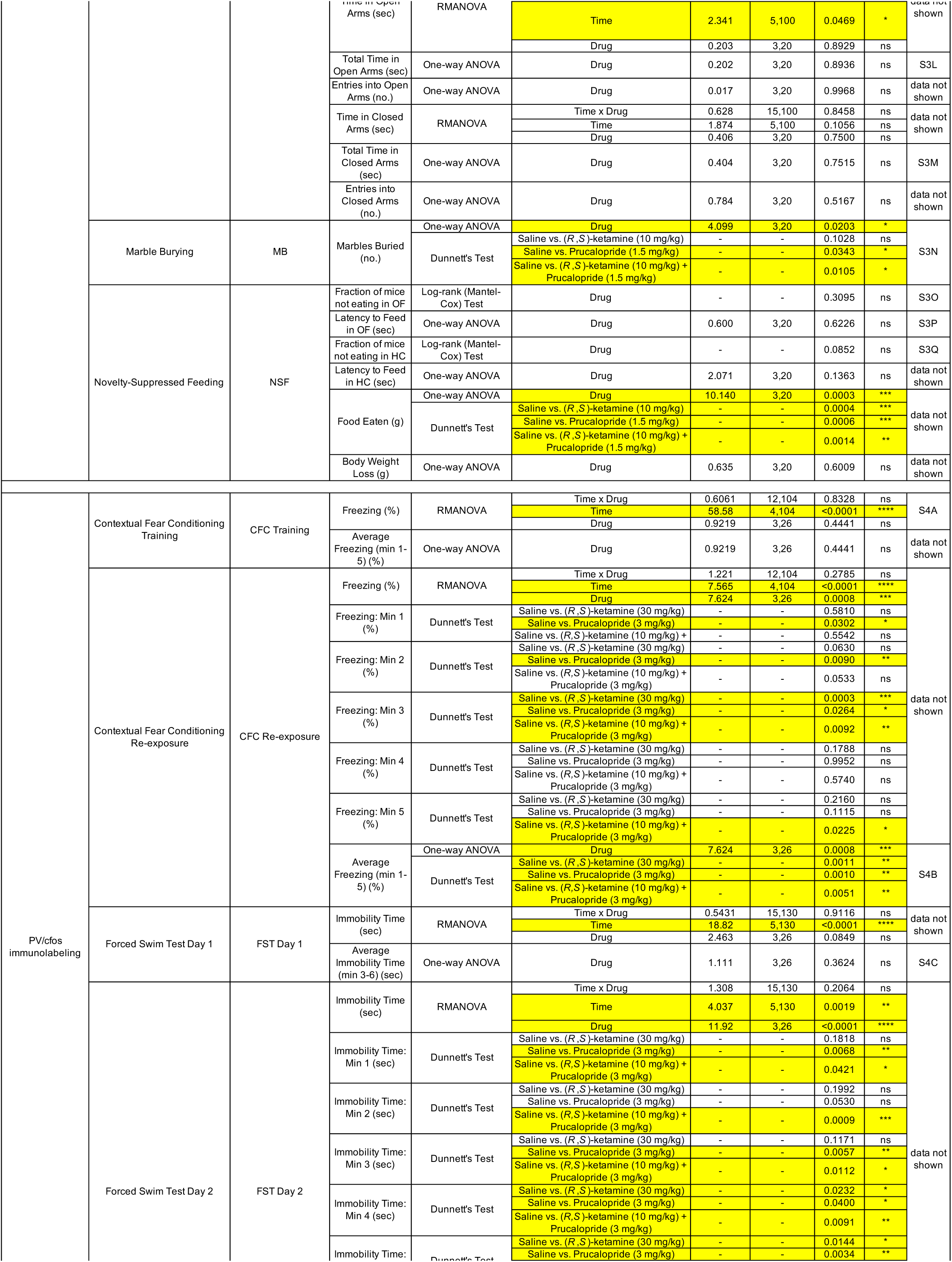
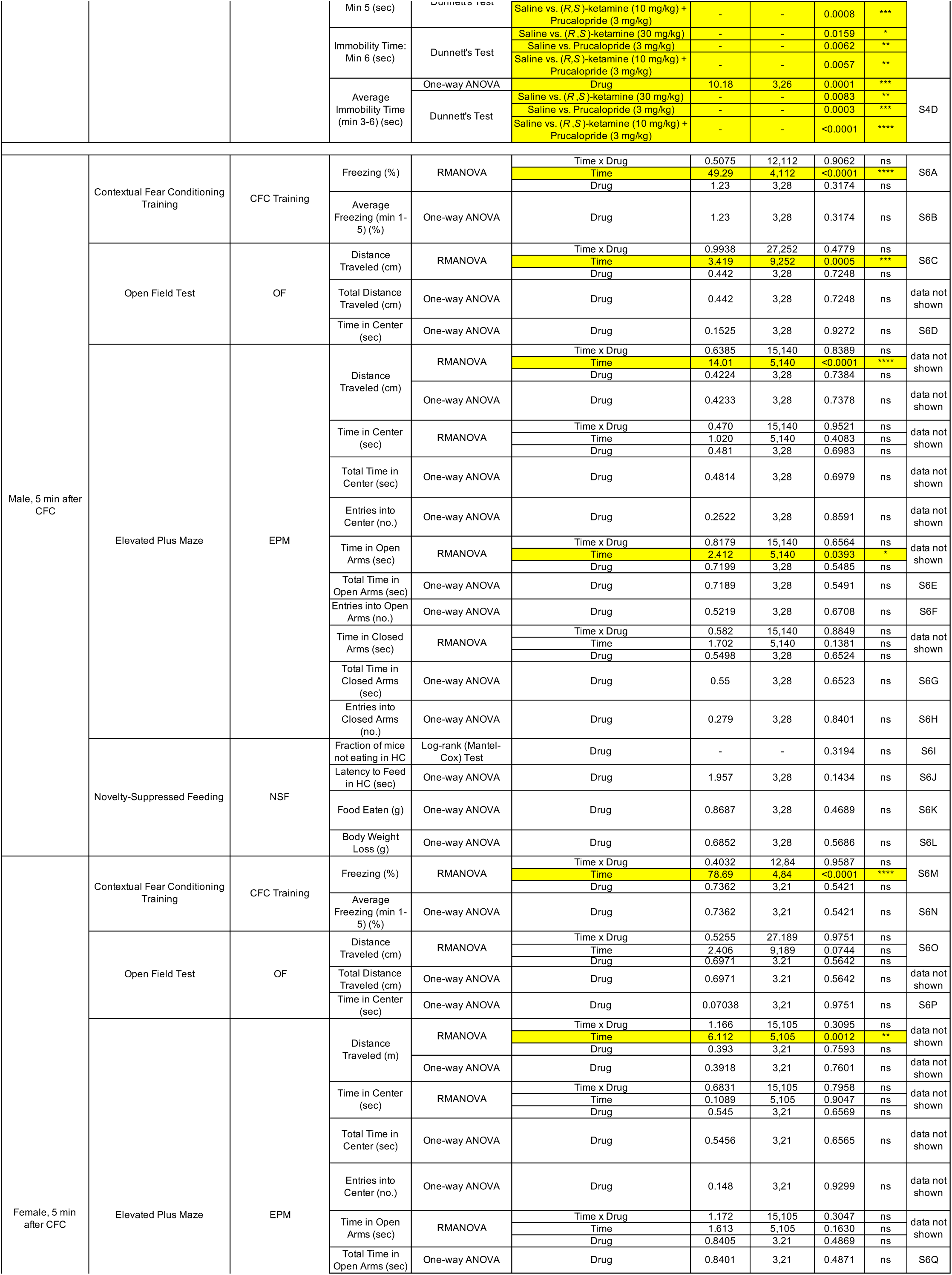
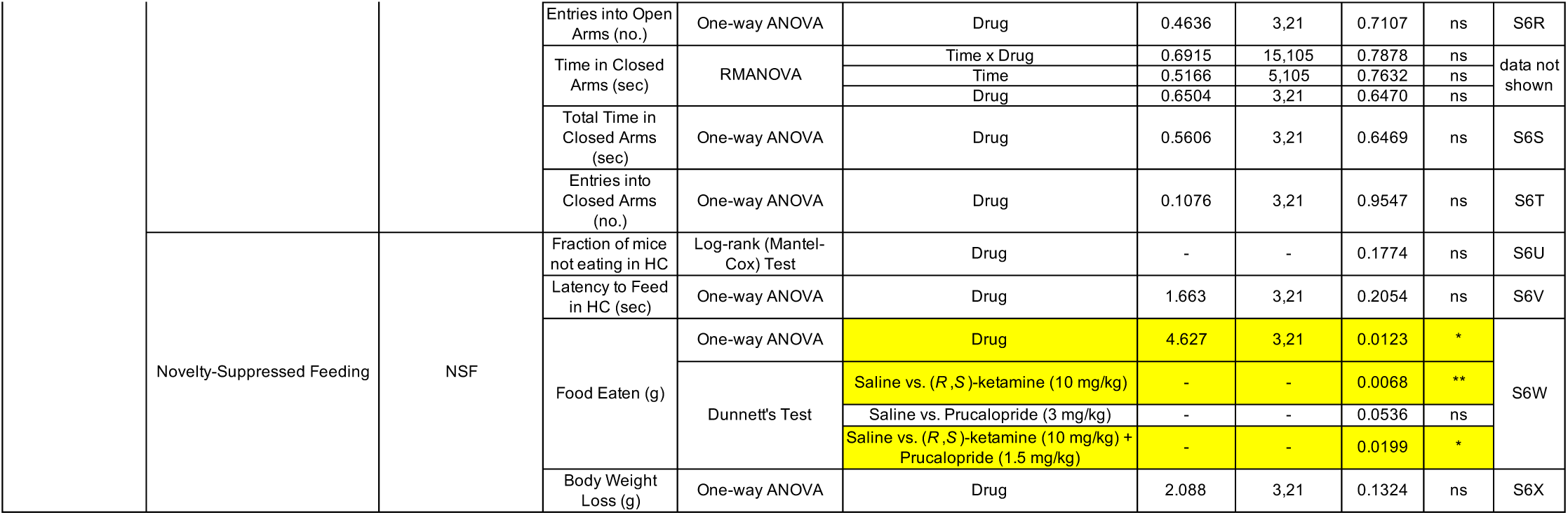
Behavioral and electrophysiological statistical analysis.

**Table S02.**
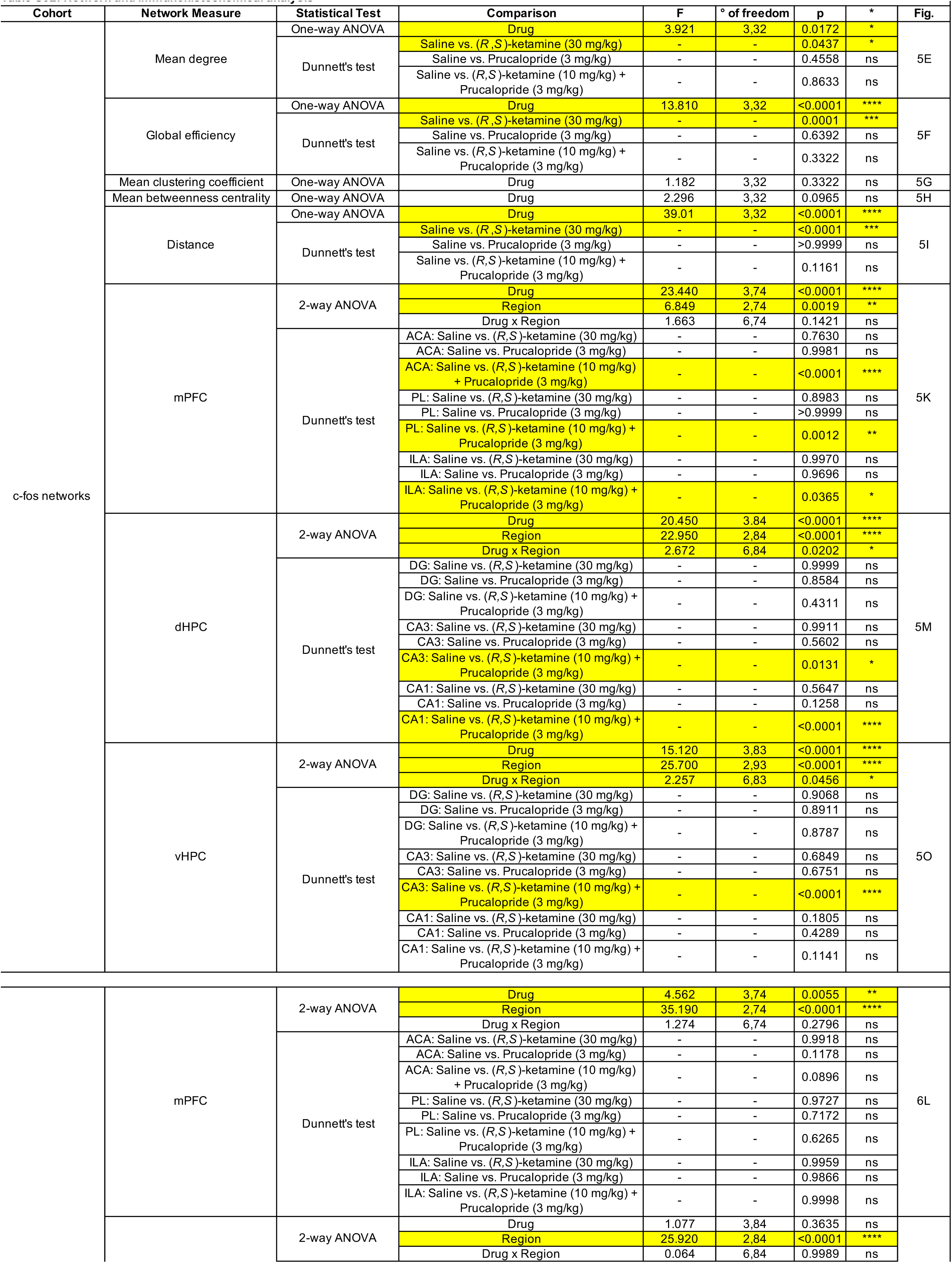
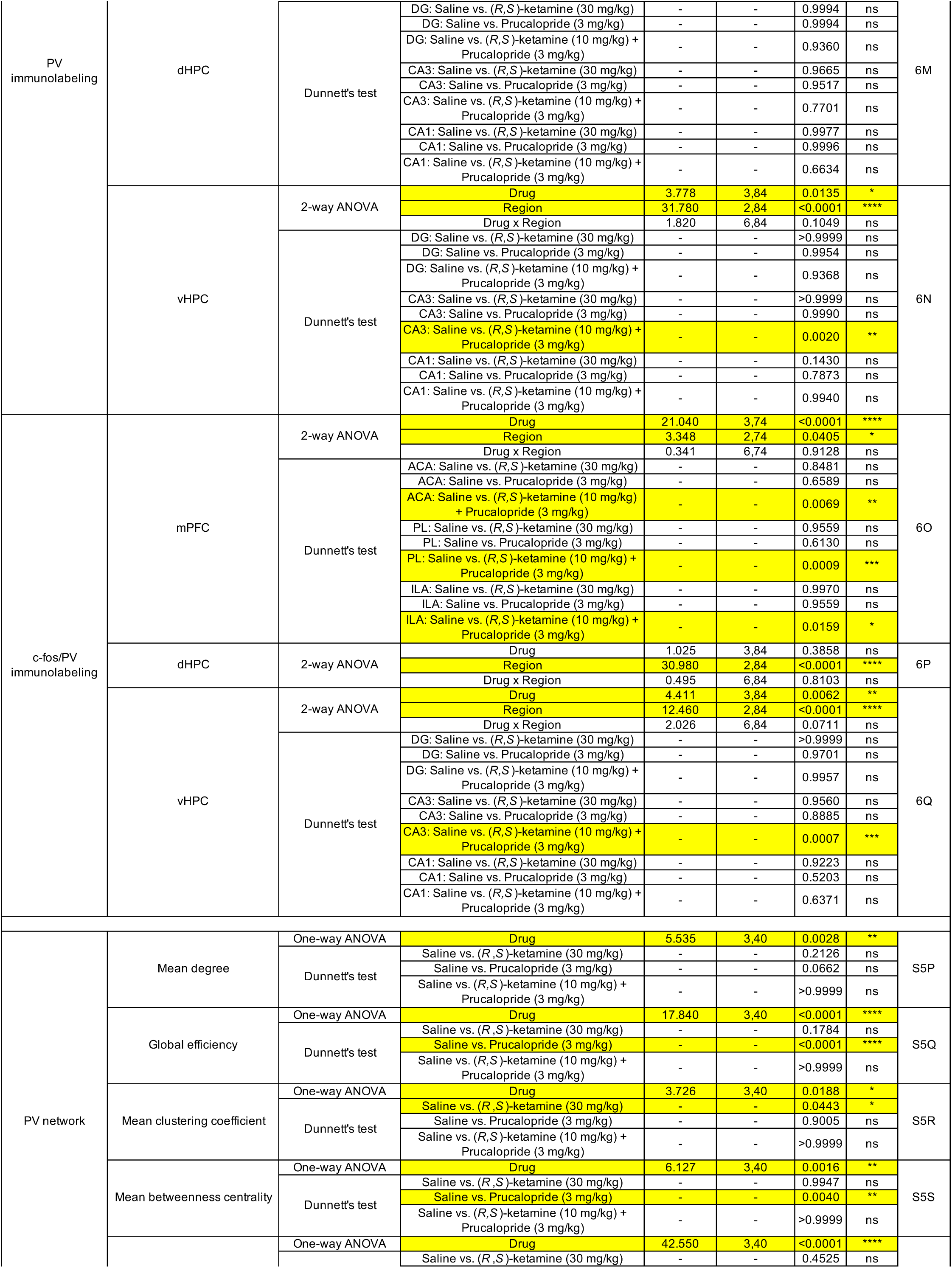
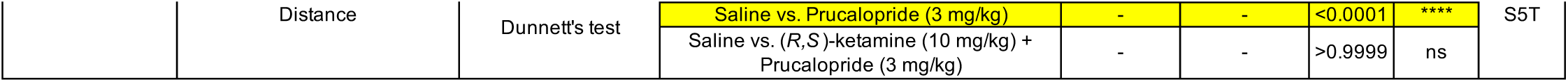
Network and immunohistochemical analysis.

**Table S03.**
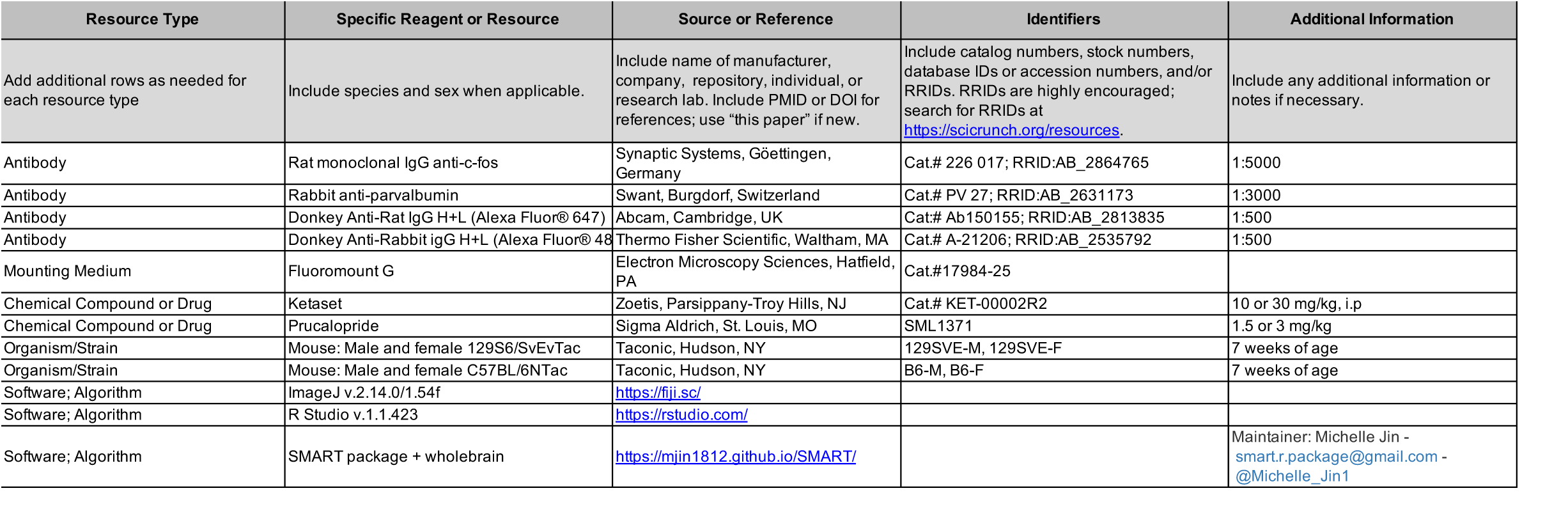
Key resources.

